# The *Nkx2.3*–*Nr5a1* gene cascade plays a crucial role in spleen-specific vascular architecture and marginal zone formation

**DOI:** 10.64898/2026.02.02.703419

**Authors:** Kanako Miyabayashi, Koji Ono, Tetsuya Sato, Ayano Yahagi, Masanori Iseki, Katsuhiko Ishihara, Takami Mori, Miki Inoue, Ryuki Shimada, Kei-ichiro Ishiguro, Tomohiro Ishii, Jongsung Noh, Man Ho Choi, Takashi Baba, Yasuyuki Ohkawa, Emi Kiyokage, Kazunori Toida, Yuichi Shima

## Abstract

NR5A1 is a nuclear receptor and master regulator of steroidogenic tissue development and steroid hormone biosynthesis in the adrenal glands and gonads. Although NR5A1 is expressed in splenic vascular endothelial cells, its function in the adult spleen remains unclear. Using single-cell transcriptomics in mice, we show that NR5A1-positive endothelial cells are heterogeneous and exhibit distinct phenotypes according to their localization in the splenic sinus or marginal sinus. To assess NR5A1 function, we generated a spleen-specific *Nr5a1* enhancer deletion model. Enhancer loss resulted in reduced spleen size and ablation of the marginal zone and marginal sinus. In addition, 21α-hydroxylase activity in splenic sinusoidal endothelial cells was reduced, particularly in females, leading to decreased levels of adrenocortical steroids in the spleen. Enhancer deletion also induced ectopic lymphatic-like and high endothelial venule-like vasculature similar to those observed in *Nkx2.3*-deficient mice. Consistently, mutation of the NKX2.3-binding motif within the enhancer abolished NR5A1 expression. Together, these findings identify an *Nkx2.3*→*Nr5a1* regulatory axis essential for spleen-specific endothelial cell differentiation and splenic architecture required for blood filtration and marginal zone formation.

## Introduction

The spleen is a central secondary lymphoid organ that performs two key physiological functions. It maintains erythrocyte homeostasis by clearing senescent red blood cells and recycling iron, and it serves as a critical site for immune surveillance and immune responses against blood-borne antigens. The spleen comprises two functionally distinct compartments: red and white pulp. Blood from the splenic artery enters the central artery, which is sheathed by lymphocytes to form the white pulp. A portion of the arterial blood flows into the marginal sinus, located at the interface between the white and red pulp, while the remainder enters the red pulp. In the red pulp, erythrocytes released into open circulation are reclaimed into the large sinusoidal lumen through slits between rod-shaped endothelial cells ^1^. Thus, the red pulp serves as a blood filter, where aged or inflexible erythrocytes are trapped and phagocytosed by macrophages^2^. In the white pulp, T cells form periarterial lymphoid sheaths, whereas B cells establish lymphoid follicles that support clonal expansion and immunoglobulin production. The venous sinus surrounding the white pulp, termed the marginal sinus, serves as an entry site for immune cells to migrate into the white pulp^3^. Various subsets of immune cells, including SIGNR1 (CD209b)-positive macrophages, MARCO-positive marginal zone macrophages^4,5^, CD169 (SIGLEC-1/sialoadhesin)-positive marginal zone metallophilic macrophages^6,7^, and marginal zone B cells^8^, reside in the marginal zone and maintain the functional and structural integrity of this unique histological region. Marginal-zone macrophages recognize and phagocytose blood-borne antigens. Marginal-zone B cells rapidly respond to blood-borne antigens by producing IgM and playing distinct roles in follicular B cells, which initiate adaptive responses by undergoing class-switch recombination and producing high-affinity antibodies^9–11^. Recent studies have shown that CD169-positive macrophages migrate to myocardial infarction lesions and contribute to cardiac recovery by suppressing inflammation and regulating fibrosis, highlighting the novel physiological role of marginal-zone macrophages^12^.

NR5A1 (also known as Ad4BP or steroidogenic factor-1) is a nuclear receptor critical for the development of steroidogenic tissues, such as the adrenal glands and gonads^13^. NR5A1 is expressed in splenic sinusoidal endothelial cells, and mice with *Nr5a1* gene disruption display a marked reduction in spleen size during fetal stages^14^, suggesting impaired splenic blood filtering function. However, because these mice die shortly after birth due to adrenal insufficiency^15^, the role of NR5A1 in the adult spleen remains unclear. Recent single-cell transcriptomic analyses of vascular endothelial cells across various mouse tissues have revealed that genes involved in steroid hormone biosynthesis are expressed in splenic endothelial cells^16^. Although NR5A1 is known to regulate the expression of steroidogenic enzymes in the adrenal cortex^17^, the relationship between NR5A1 and steroid hormone production in splenic endothelial cells remains unclear. Furthermore, the physiological significance of steroid hormone production in the spleen remains to be elucidated.

NKX2.3 is a homeobox transcription factor that plays a crucial role in splenic venous sinus formation. *Nkx2.3* (*Nkx2-3*) knockout mice exhibit perinatal lethality and display defects in the spleen and gut-associated lymphoid tissues^18^. In particular, the marginal zone is severely disrupted in *Nkx2.3*-deficient spleens^19,20^, and lymphoid cysts filled with lymphocytes and high endothelial venules (HEVs)—structures normally restricted to lymph nodes—develop in knockout spleens^21,22^. These findings establish *Nkx2.3* as essential for spleen-specific vascular architecture.

In this study, we used single-cell transcriptomic analyses to demonstrate that NR5A1-positive endothelial cells in the spleen exhibit distinct gene expression profiles in the splenic and marginal sinuses. We further identify a spleen-specific enhancer that governs *Nr5a1* expression in endothelial cells of both the splenic and marginal sinuses. Through genetic perturbation, we demonstrate that endothelial NR5A1 function is indispensable for establishing spleen-specific vascular identity and marginal zone formation. Furthermore, we identify NKX2.3 as a direct upstream regulator of this enhancer, defining a conserved transcriptional axis that links vascular specialization to immune organ architecture.

## Materials and Methods

### Mouse strains

Ad4BP-BAC-EGFP mice^23^ were used to visualize the cytoplasm of the NR5A1-expressing sinus endothelium and to collect NR5A1-positive endothelial cells from the spleen by cell sorting. StAR/eGFP mice have been described previously^24^. *Nr5a1^ΔSpE^*mice were generated using genome editing^25–27^. Guide RNAs targeting the upstream and downstream regions of the SpE were designed using CRISPR Direct (http://crispr.dbcls.jp/), and crRNA, tracrRNA, and Cas9 protein (Integrated DNA Technologies) were mixed and introduced into the fertilized eggs by electroporation (Genome Editor, BEX). For the generation of *Nr5a1^SpE-R2m^* mice, guide RNA targeting the R2 sequence in the SpE and donor DNA containing a mutated R2 sequence (R2m) were used. The eggs were then transferred into the oviducts of recipient mothers, and the genotypes of the pups were analyzed by PCR. Integration of the R2m sequence was confirmed by *Bam*HI digestion of the PCR-amplified DNA fragments. Sequences of the guide RNAs and genotyping primers are listed in Supplementary Table 1. All animal protocols were approved by the Animal Care and Use Committee of Kawasaki Medical School and Kurume University School of Medicine.

### Single-cell transcriptome analyses of NR5A1^+^ endothelial cells

Ad4BP-BAC-EGFP mice were sacrificed at approximately P70, and their spleens were collected (n = 3, 2 males and 1 female). Spleens were mechanically minced and enzymatically digested with collagenase (Nacalai Tesque). The isolated cells were subjected to fluorescence-activated cell sorting (FACS) using BD FACSAria III (BD) or CytoFLEX SRT (Agilent Biotechnologies). EGFP-positive cells corresponding to NR5A1-expressing vascular endothelial cells were subjected to single-cell RNA sequencing (scRNA-seq) library preparation using the 10x Genomics Chromium Single Cell 3′ Library & Gel Bead Kit v3.1, following the manufacturer’s protocol. Libraries from three independent batches were sequenced on an Illumina NovaSeq 6000 platform, and the resulting FASTQ files were processed using Cell Ranger (v9.0.0) with default settings. Reads were aligned to a custom reference genome constructed by appending the enhanced green fluorescent protein (EGFP) gene sequence to the mouse genome assembly (mm10). Initial data quality control and generation of feature-barcode matrices were performed using Cell Ranger’s “count” pipeline. Subsequent analyses were performed in R using the Seurat package (v5.1.0)^28^. Cells with fewer than 500 detected genes, more than 6,000 unique molecular identifiers (UMIs), or more than 10% of reads mapping to mitochondrial genes were excluded to eliminate low-quality cells and potential doublets. Potential doublets were further removed using DoubletFinder (v2.0.6)^29^, with the doublet formation rate estimated at 7.5% and the optimal pK value determined using the “find.pK” function. After quality filtering was applied, including the EGFP expression criterion, 31,199 high-quality cells were retained across the three batches. Gene expression data were normalized using the “NormalizeData” function, and batch effects across samples were simultaneously corrected and integrated using the Harmony algorithm^30^. Dimensionality reduction was performed using principal component analysis (PCA), and the top 30 principal components were used for clustering via the Louvain algorithm at a resolution of 0.1 using the “FindClusters” function. Clusters were visualized using uniform manifold approximation and projection (UMAP) dimensionality reduction implemented via the “RunUMAP” function. From the above analyses, six cell clusters were identified. Differentially expressed genes (DEGs) between clusters were identified using the “FindAllMarkers” function employing the Wilcoxon rank-sum test with the Bonferroni correction for multiple comparisons. Marker genes were defined as those with an average log2 fold change ≥ 1 and an adjusted p-value < 0.01. To functionally annotate the marker genes in each cluster, Gene Ontology (GO) enrichment analysis was performed using the R package clusterProfiler (v 4.17.0)^31^. For each cluster, the top 150 DEGs were selected based on descending log2 fold-change values. These genes were then analyzed using the “compareCluster” and “enrichGO” functions with a q-value cutoff set to 0.01. The analysis was conducted across three GO categories: biological processes (BP), molecular functions (MF), and cellular components (CC).

### Western blot analyses

Nuclear extracts were prepared from adrenal glands and spleens (Nuclear Extraction Kit, ab113474, Abcam) and quantified using a Qubit Protein Assay (Thermo Fisher Scientific). Forty micrograms of protein was subjected to SDS-polyacrylamide gel electrophoresis and western blot analyses. Primary and secondary antibodies used in the analyses are listed in Supplemental Table 2.

### Tissue preparation, histological studies, and immunostaining

Mice were anesthetized with 0.3 mg/kg medetomidine hydrochloride (Nippon Zenyaku Kogyo), 4 mg/kg midazolam (Astellas Pharma), and 5 mg/kg butorphanol tartrate (Meiji Seika Pharma). Fresh samples were isolated, and bright-field or fluorescence images of the specimens were acquired using a stereomicroscope (Evident SZx10 or SZX16). For histological analyses, cardiac perfusion fixation was performed using PBS, followed by 4% paraformaldehyde in PBS. The isolated tissues were fixed by immersion in 4% paraformaldehyde in PBS and subjected to subsequent analyses.

The tissues were embedded in paraffin wax, sectioned at 5-μm thickness, and subjected to hematoxylin and eosin (H&E) and elastica van Gieson (EVG) staining. Images were obtained using BZ-X700 (Keyence), and H&E images were used to measure the area of white pulp and the total area of the spleen using the MZ-X Analyzer 1.3.0.3 (Keyence). For immunostaining, samples were cut into 50-μm-thick sections using a CM3050 S cryotome (Leica). The sections were stained by the free-floating staining method^27^. Primary and secondary antibodies used in this study are listed in Supplementary Table 2. 4′6′-Diamidino-2-phenylindole (DAPI) (Sigma) was used for nuclear staining. Tissue sections were encapsulated in VECTASHIELD Mounting Medium (Vector Laboratories) and photographed using a BZ-X700 (Keyence) or LSM 700 laser-scanning microscope (Zeiss).

### Analyses of the circulating hemocytes

Blood smear samples were stained with Giemsa, and the number of erythrocytes containing Howell–Jolly bodies was determined. The erythrocytes were counted using an Olympus Motorized Microscope System BX61 equipped with a DP74 digital camera (Olympus). At least 1,000 erythrocytes were counted in each specimen (n = 4 for each experimental group).

### Scanning electron microscopic (SEM) studies

Spleens were fixed by perfusion fixation, followed by interstitial perfusion^32^ and immersion fixation using 2.5% glutaraldehyde (TAAB) and 4% paraformaldehyde (Merck) in 0.1 M phosphate buffer (pH 7.4). Tissue blocks (4 × 4 × 2 mm) were trimmed from the fixed organs. After post-fixation with 1% OsO_4_, the buffer was replaced with 50% DMSO. Specimens immersed in 50% DMSO were quench-frozen in liquid nitrogen and fractured by mechanical impact. Fixation with 1% OsO_4_ was repeated. Then, the specimens were immersed in 2% tannic acid for 16 h and in 2% OsO_4_ for 6 h according to the previously reported “tannin-osmium method”^33^. Next, the specimens were dehydrated in increasing concentrations of ethanol and transferred to *t*-butylalcohol, and the soaked samples were sublimed in vacuum and dried while frozen. The specimens were attached, with the fractured surface facing upwards, on a metal stub with a conductive paste. All the specimens were lightly coated with osmium (Neoc-Pro Osmium Coater, Meiwafosis) and platinum (JEC-3000FC Auto Fine Coater, JEOL) by vacuum evaporation. The specimens were observed using an S-3400N scanning electron microscope (Hitachi) at an accelerating voltage of 15 kV.

### Bulk mRNA-sequencing, data processing, and differential gene expression analyses

mRNA sequences were analyzed as previously described^26,27^. Briefly, total RNA was prepared from spleens or sorted endothelial cells and subjected to library construction using the Illumina TruSeq Library Construction Kit (Illumina) or NEBNext Ultra RNA Library Prep Kit (New England Biolabs). Libraries were sequenced using a NovaSeq 6000 sequencer (Illumina) with 150-bp paired-end reads. After adapter sequences and low-quality sequences were removed using Cutadapt with default parameters, FASTQ files were aligned to the mouse genome (mm10) using STAR (v2.7.3a) with default parameters. FeatureCounts (v2.0.1) and edgeR (v3.28.1) were used to assemble the transcripts and calculate the counts per million (CPM) of each gene using default parameters. DEGs were identified using the criteria of |Log2FC | ≥ 2, *p* < 0.01, and FDR < 0.01, and were subjected to GO analyses using Metascape^34^ (https://metascape.org/gp/index.html#/main/step1).

### Flow cytometry analyses

Flow cytometry analysis of immune cells in the spleen was performed as previously described^35^. The antibodies used for the flow cytometry analyses are listed in Supplemental Table 2.

### Measurement of steroid hormones in the spleen

Spleens were collected from the control and ΔSpE mice approximately 10 weeks after birth (n = 3 per experimental group). Steroid levels were determined by liquid chromatography-mass spectrometry (LC-MS) following established methods^36,37^ and are expressed as nanograms per gram of tissue. The metabolic ratio for each enzymatic reaction was calculated by dividing the metabolite concentration by the precursor concentration.

### Sequence data deposition in a public database

All sequencing data were deposited in the Gene Expression Omnibus (GEO) as follows: bulk RNA-seq data of the spleen: GSE303957, bulk RNA-seq data for NR5A1-positive cells isolated by sorting: GSE303956, and scRNA-seq data for NR5A1-positive cells isolated by sorting: GSE303958.

### Statistical analyses

At least three animals were used for each experimental group, unless otherwise specified. Data are presented as the mean ± SEM. Statistical comparisons between two groups were performed using the two-tailed Student’s *t*-test, and comparisons among multiple groups were conducted using one-way ANOVA followed by the Tukey–Kramer post hoc test; *p* < 0.05 was considered statistically significant.

## Results

### Spatial heterogeneity of NR5A1-expressing endothelial cells in the spleen

NR5A1 has been shown to be expressed in the vascular endothelial cells of the spleen^14^. However, its expression in the spleens of adult mice has not been thoroughly investigated. To address this, we analyzed EGFP expression in the spleens of Ad4BP-BAC-EGFP transgenic mice^23^, which faithfully recapitulate endogenous NR5A1 expression. EGFP expression was macroscopically heterogeneous within the spleen and was particularly prominent in the red pulp (Fig. 1A, 1A’). Immunostaining confirmed that EGFP almost completely overlapped with endogenous NR5A1 in the spleen (Fig. S1). Immunostaining of EGFP with the T and B cell markers CD3ε and CD45R, respectively, indicated that EGFP was predominantly expressed in the endothelial cells of the splenic sinusoids within the red pulp (Fig. 1B, 1C). To investigate the location of EGFP-positive cells in more detail, EGFP was compared with several marker proteins for macrophage subtypes (F4/80, macrophage receptor with a collagenous structure [MARCO], and CD169) and a marker for marginal sinus-lining cells (mucosal vascular addressin cell adhesion molecule-1 [MAdCAM-1]). EGFP-positive cells were distributed in the red pulp, where F4/80-positive macrophages are present (arrows in Fig. 1D, D’, D’’). EGFP-positive cells were also found in the marginal zone, where MARCO-positive macrophages reside (arrowheads in Fig. 1E, E’, E’’). CD169-positive macrophages (also known as metallophilic macrophages) were located on both the white pulp and red pulp sides of the marginal sinus^7^, suggesting that EGFP was expressed in the endothelial cells of the marginal sinus (arrowheads in Fig. 1F, F’, F’’). MAdCAM-1 has been reported to mark the “lining cells” of the marginal sinus^38^, and our analyses also found that MAdCAM-1 is expressed in thin cells, which are closely associated with EGFP-positive cells (arrowheads in Fig. 1G, G’, G’’) (Fig. S2). Collectively, these data confirm that EGFP is expressed in the endothelial cells of the marginal sinus. Figure 1H summarizes the schematic representation of spatial distribution of NR5A1-positive cells and other cell types in the marginal sinus and marginal zone.

**Figure 1.**
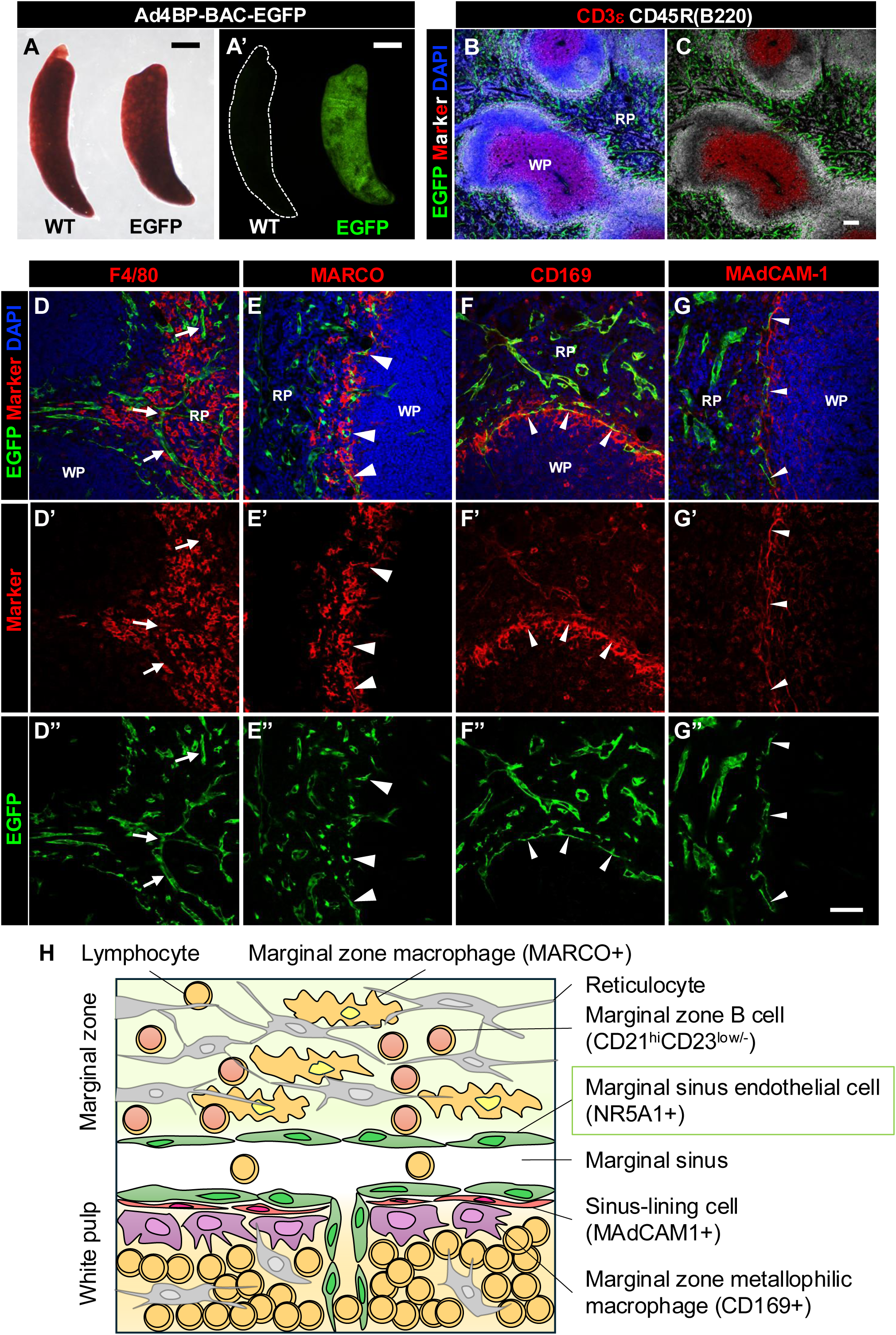
EGFP expression in the spleen of Ad4BP-BAC-EGFP mice. (A, A’) Macroscopic view of wild-type and Ad4BP-BAC-EGFP mouse spleens under bright-field (A) and EGFP fluorescence (A’) microscopy. The wild-type spleen is outlined with a white dotted line in A’. Bar = 500 µm. (B, C) Immunostaining for EGFP together with CD3ε and CD45R in Ad4BP-BAC-EGFP mouse spleens, shown with (B) or without (C) DAPI nuclear counterstaining. Bar = 100 µm. (D-D’’, E-E’’, F-F’’, G-G’’) Immunostaining of EGFP with cell type-specific markers: F4/80 (D-D’’), MARCO (E-E’’), CD169 (F-F’’), and MAdCAM-1 (G-G’’). Arrows indicate EGFP expression in the endothelial cells of the venous sinuses in the red pulp (splenic sinuses), and arrowheads indicate EGFP expression in the endothelial cells of the venous sinuses at the border between the white pulp and the marginal zone (marginal sinuses). Bar = 50 µm. (H) Schematic illustration of the histological structure of the marginal zone and sinus, showing the cell types observed in this region. RP, red pulp; WP, white pulp.

### Functional differences between endothelial cells of the splenic sinus and marginal sinus

scRNA-seq analysis of EGFP-positive cells sorted from the spleens of Ad4BP-BAC-EGFP mice was performed. The dimensional reduction of gene expression and clustering of cells collected from three distinct mice (two males and a female) showed almost the same pattern (Fig. S3A), comprising six clusters, including two major ones (clusters 1 and 2 in Fig. 2A). EGFP and endogenous NR5A1 expression was detected in all clusters (Fig. 2B and 2C). Moreover, the sinusoidal endothelial marker *Stab2* was detected across all clusters, including two major clusters (Fig. 2D), confirming the validity of our scRNA-seq analysis. To clarify the relationship between the scRNA-seq results and the spatial distribution of NR5A1-expressing cells observed by immunostaining, we focused on *Star*, which was highly expressed in cluster 1 (Fig. 2E), and *Cd34*, which was highly expressed in cluster 2 (Fig. 2F). Immunostaining using Ad4BP-BAC-EGFP mice^23^ revealed that CD34 was expressed in the endothelial cells of the marginal sinus (arrows in Fig. 2G). Furthermore, using StAR/eGFP reporter mice^24^, in which EGFP expression recapitulates endogenous StAR expression, we observed that CD34-positive and StAR-positive cells were largely mutually exclusive, although a subset of cells co-expressed both markers (arrows in Fig. 2H). Notably, StAR-positive endothelial cells were located outside the marginal zone, where MARCO-positive macrophages are found (Fig. 2I), and within the red pulp, where F4/80-positive macrophages are found (Fig. 2J), strongly suggesting that these cells represent the endothelial cells of the splenic sinus. Thus, cluster 1 in scRNA-seq corresponded to the endothelial cells in the splenic sinus, whereas cluster 2 represented the endothelial cells of the marginal sinus.

**Figure 2.**
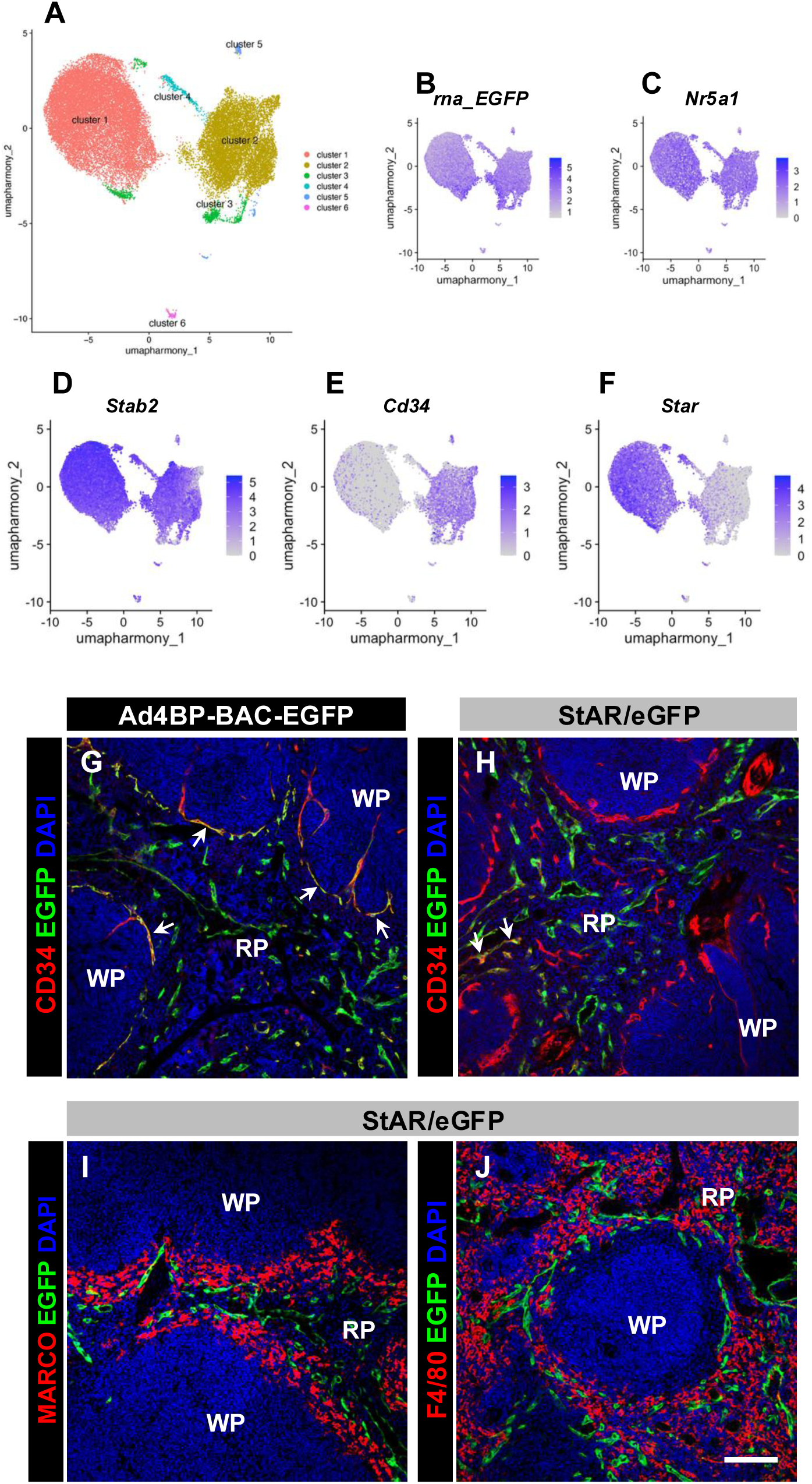
Functional and spatial heterogeneity of EGFP-positive endothelial cells in the spleens of Ad4BP-BAC-EGFP mice. (A) UMAP plot of clusters consisting of EGFP-positive endothelial cells isolated from Ad4BP-BAC-EGFP spleens. (B–F) Feature plots of representative genes expressed in EGFP-positive endothelial cells: *Egfp* (B), *Nr5a1* (C), *Stab2* (D), *Cd34* (E), and *Star* (F). (G) Immunostaining of EGFP and CD34 in Ad4BP-BAC-EGFP spleens. (H–J) Immunostaining of EGFP with CD34 (H), MARCO (I), or F4/80 (J) in StAR/eGFP spleens. Arrows in (G) indicate CD34/EGFP double-positive cells in Ad4BP-BAC-EGFP spleens, whereas arrows in (H) indicate CD34/EGFP double-positive cells in StAR/eGFP spleens. Bar = 100 µm. RP, red pulp; WP, white pulp.

To elucidate the differences in the properties of endothelial cells between the splenic and marginal sinuses, we performed a more detailed analysis of gene expression in each cluster. Markers commonly expressed in endothelial cells—*Pecam1*, *Cdh5*, *Eng*, and *Erg*—were detected in all clusters, albeit at varying expression levels (Fig. S3B). Gene Ontology (GO) analysis was performed using the top 150 highly expressed genes in each cluster (Fig. 3A, Fig. S4; the gene list is shown in Supplemental Table 3). Cluster 1 was characterized by genes associated with negative chemotaxis (*Flrt2*, *Sema3*d, *Itga9*, *Mmp28*, *Robo2*) and extracellular matrix organization (*Adamts5*, *Col23a1*) (Fig. 3B and Fig. S3C). These genes are also associated with the induction of vasculogenesis, tissue remodeling, and angiogenesis guidance. *Flrt2*, *Sema3d*, and *Robo2* are guidance molecules, and *Jag1* encodes a Notch ligand involved in endothelial fate determination. *Gli3* encodes a transcription factor in the Hedgehog signaling pathway. *Ccbe1*, *Itga9*, *Mmp28*, in addition to *Adamts5* and *Col23a1*, are involved in extracellular matrix degradation and remodeling^39^. In contrast, genes highly expressed in cluster 2 were associated with GO terms, such as endothelial cell migration (*Ptp4a3*, *Dll4*, *Col18a1*, *Kdr*, *Id1*, *Adamts9*, *Amotl2*) and angiogenesis (*Esm1*, *Dll4*, *Mcam*, *Emcn*) (Fig. 3A, 3C, Fig. S3D). These terms suggest active angiogenesis. Notably, *Esm1*, *Dll4*, *Kdr*, and *Id1* are known markers of tip cells^40–43^. Moreover, *Esam*, *Col18a1*, *Adamts9*, and *Mcam* have been implicated in cell–cell or cell–matrix adhesion, further pointing to their relevance to angiogenesis^44^.

**Figure 3.**
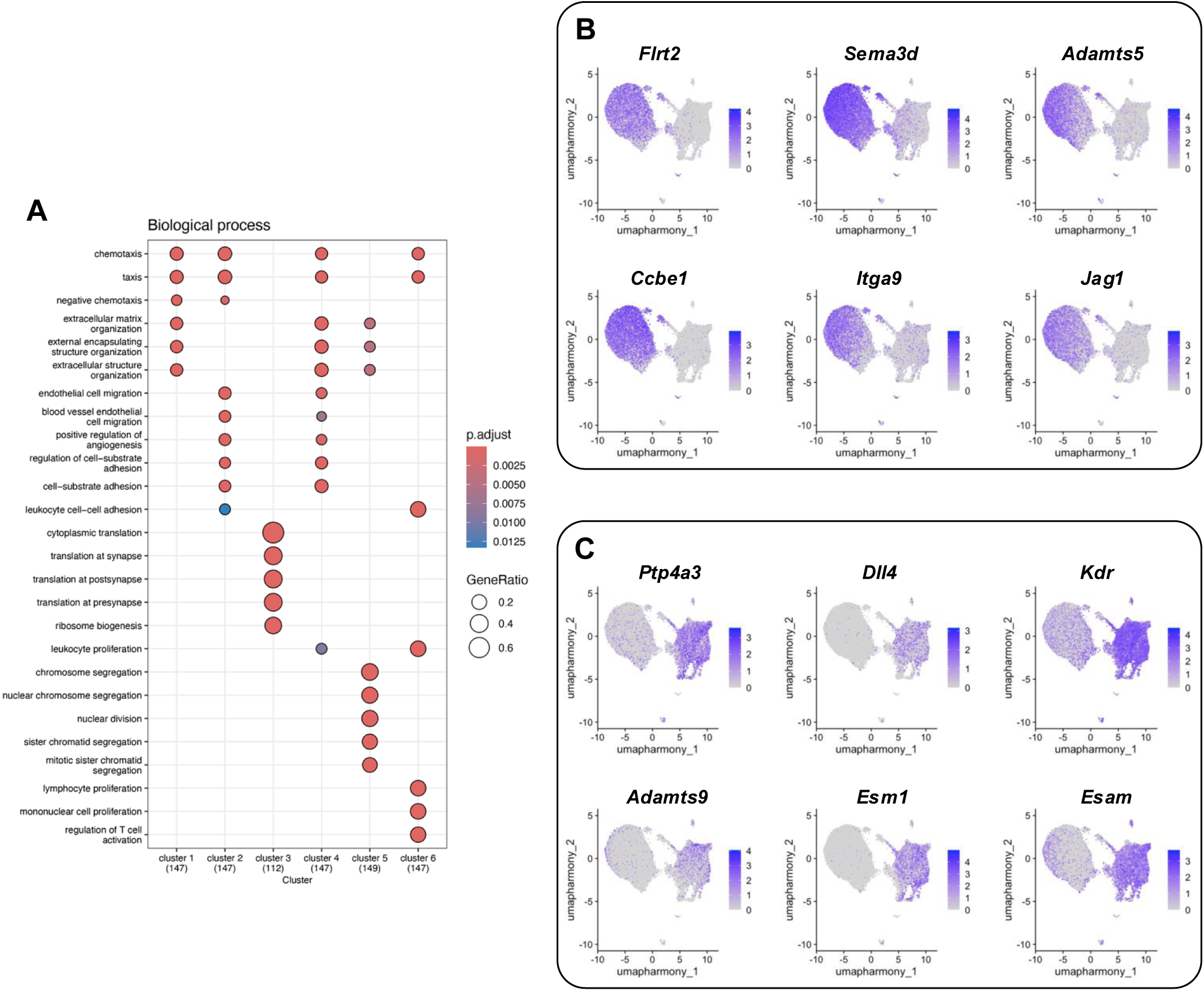
Functional characterization of clusters constituting of EGFP-positive endothelial cells. (A) Bubble plot showing the biological processes characterizing each cluster. The top 150 highly expressed genes in each cluster were used for GO analysis. (B) Feature plots of representative genes highly expressed in cluster 1: *Flrt2*, *Sema3d*, *Adamts5*, *Ccbe1*, *Itga9*, and *Jag1*. (C) Feature plots of representative genes highly expressed in cluster 2: *Ptp4a3*, *Dll4*, *Kdr*, *Adamts9*, *Esm1*, and *Esam*.

Collectively, these results revealed that cells in cluster 1 (splenic sinus) highly express genes involved in vascular patterning and guidance, whereas cells in cluster 2 (marginal sinus) exhibit high expression of genes related to active angiogenesis. Cluster 3 exhibited a high abundance of ribosomal genes (Fig. 3A, Fig. S4A, S4B), suggesting that it represents a population of dead or low-quality cells. Although the developmental relationship between clusters 1 and 2 remains unclear, cluster 4 exhibited an intermediate gene expression profile, suggesting a transitional state between the two major clusters (Fig. 3A, Fig. S4A, S4B). Cluster 5 was characterized by genes related to chromosomal segregation, nuclear division, and sister chromatid segregation, suggesting that these cells were mitotic (Fig. 3A, S4A, S4B). Cluster 6 was enriched in genes related to neutrophil (*Ikzf1*) and lymphocyte differentiation (*Igkc*, *Ly6d*) (Fig. 3A, Fig. S3E, Fig. S4A, S4B), suggesting that this cluster consisted of doublets or contaminating cells, including neutrophils and lymphocytes.

### Identification of the SpE in the Nr5a1 gene

We previously identified several tissue-specific enhancers in the *Nr5a1* locus, including the ventromedial hypothalamic nucleus enhancer (VE)^45^, fetal adrenal gland enhancer (FAdE)^46^, pituitary gonadotrope enhancer (PE)^47^, and fetal Leydig cell enhancer (FLE)^48^ (Fig. 4A).

**Figure 4.**
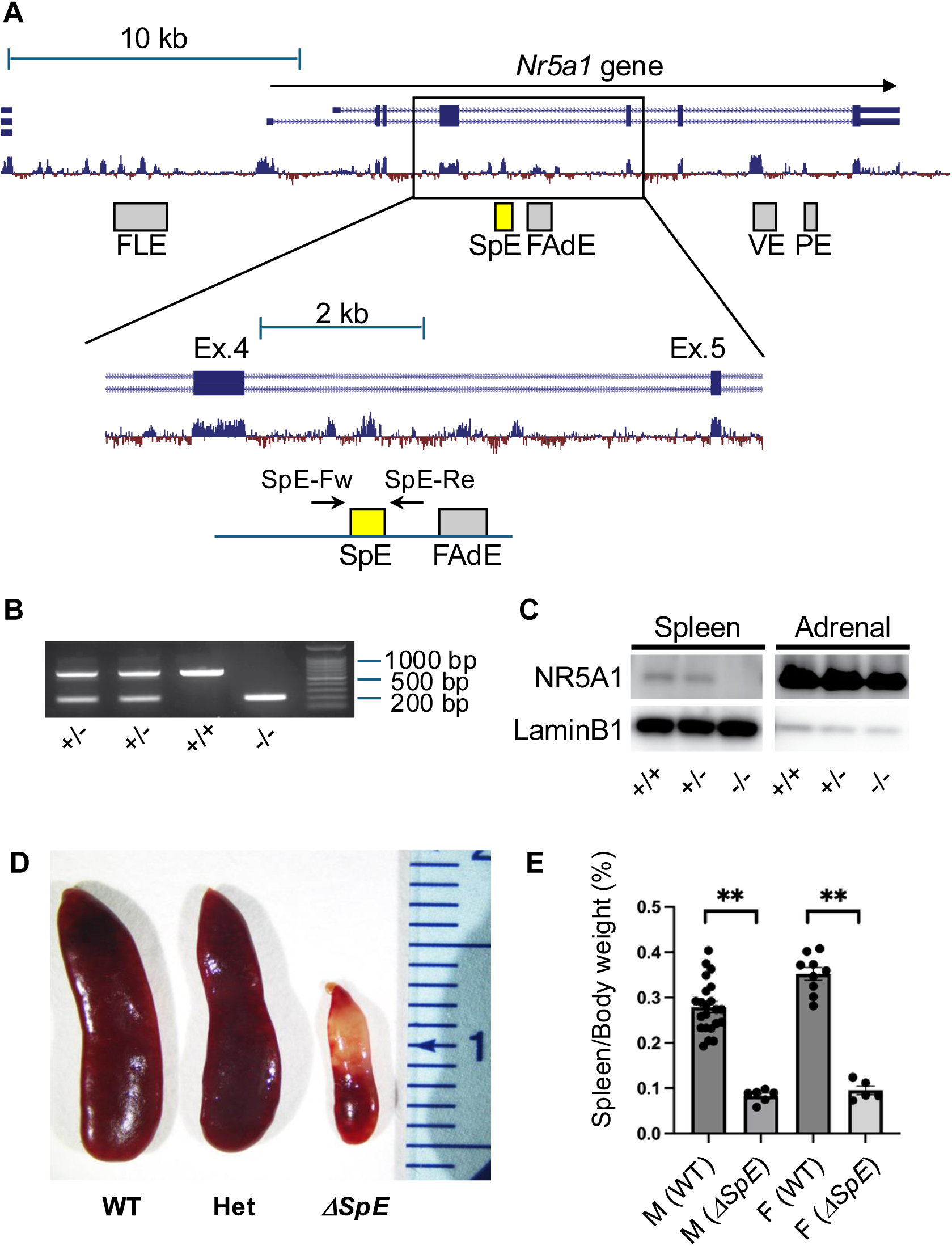
Spleen-specific depletion of NR5A1 by deletion of the spleen enhancer (SpE) of the *Nr5a1* gene. (A) Schematic representation of the *Nr5a1* gene structure and sequence conservation among mammals. Gray boxes indicate previously identified enhancers, including fetal Leydig enhancer (FLE), fetal adrenal enhancer (FAdE), ventromedial hypothalamus enhancer (VE), and pituitary enhancer (PE). The yellow box indicates the spleen enhancer (SpE) located in intron 4. Arrows (SpE-Fw and SpE-Re) denote genotyping primers. (B) PCR-based genotyping of SpE-deleted mice. The wild-type allele yields a 655-bp fragment, whereas the SpE-deleted allele yields a 213-bp fragment. (C) Loss of NR5A1 expression in the spleens of ΔSpE mice. Western blot analysis of NR5A1 and LaminB1 in nuclear extracts from spleens of wild-type (+/+), heterozygous (+/-), and homozygous (-/-) SpE-deleted mice. (D) Gross morphology of spleens from wild-type (+/+), heterozygous (+/-), and homozygous (-/-) SpE-deleted mice. (E) Spleen weight-to-body weight ratio in each genotype and sex (mean ± SE). Sample size: male wild-type, n = 12; male ΔSpE, n = 6; female wild-type n = 9; female ΔSpE, n = 5. *p* < 0.01 (**).

Furthermore, we demonstrated the functional importance of each enhancer by deleting it from the mouse genome^26,27^. We observed that the sequences of all these enhancers were highly conserved among animal species. Therefore, in this study, we focused on a small region within the fourth intron, whose sequence is conserved among mammalian species, and attempted to elucidate the *in vivo* function by deleting it using CRISPR/Cas9 genome editing (Fig. 4A). We succeeded in deleting a 441-bp region, including the conserved sequence from the mouse genome (Fig. 4B, Fig. S5). This deletion led to the complete disappearance of NR5A1 in the spleen (Fig. 4C). These results indicate that a small region in the fourth intron is indispensable for NR5A1 expression in the spleen. Therefore, we designated this region as the SpE (<u>sp</u>leen <u>e</u>nhancer) of the *Nr5a1* gene, and homozygous SpE-deletion mice were referred to as ΔSpE mice.

Macroscopically, heterozygous deletion of SpE did not markedly affect splenic size. However, the spleens of ΔSpE mice were remarkably smaller than those of the controls (Fig. 4D). The spleen, similar to the adrenal gland, exhibits sex differences in size and weight; female spleens are larger and heavier than male spleens^49^. SpE deletion affected the spleen in both males and females, resulting in a significantly lower spleen weight compared to the wild type (Fig. 4E). NR5A1 is essential for the development of the gonads and adrenal glands^15^. However, the adrenal glands, testes, and ovaries were not affected in ΔSpE mice (Fig. S6), indicating that SpE functioned only in the spleen.

### Structural and functional abnormality in ΔSpE spleens

HE staining revealed that in wild-type spleens, the white pulp was present as island-like structures within the red pulp. In contrast, in the spleens of ΔSpE mice, the white pulp in both male and female mice was fused and located centrally within the spleen, surrounded by a thin layer of red pulp (Fig. 5A). Interestingly, the measurement of the white pulp-to-red pulp ratio showed that male ΔSpE mice did not differ from wild-type mice, whereas female ΔSpE mice exhibited a significant increase compared to wild-type mice (Fig. 5B). Higher magnification of the HE-stained images of wild-type spleens showed a well-defined marginal zone between the white pulp and red pulp (Fig. 5C) and a marginal sinus at the boundary between the white pulp and the marginal zone (arrows in Fig. 5C). In contrast, in ΔSpE mice, the boundary between the white and red pulp was indistinct, and the structures of the marginal zone and marginal sinus were lost (Fig. 5D). Immunostaining analysis of wild-type spleens demonstrated the spatial arrangement patterns of B and T cells within the white pulp; CD3ε-positive T cells were localized in the central part of the white pulp, and CD45R-positive B cells surrounded the T cell area. Additionally, a marginal zone was observed outside the marginal sinus, which was stained with laminin (Fig. 5E). In contrast, the structures of the marginal zone and marginal sinus were not observed in ΔSpE mice, although the spatial distribution of B and T cells was preserved (Fig. 5F). Notably, the localization patterns of MARCO-positive macrophages in the marginal zone and CD169-positive metallophilic macrophages on the border of the white pulp and marginal sinus were disorganized in SpE heterozygous deletion mice compared with wild-type mice and were entirely lost in ΔSpE mice (Fig. 5G–I). Flow cytometry analyses demonstrated a significant decrease in CD21^hi^/CD23^low/-^ marginal zone B cells^8^ and transitional B cells of type 1 (T1B) among the B-lineage cells in ΔSpE spleens (Fig. S7); however, there were no abnormalities in the frequencies of other cell lineages in the spleen (data not shown). These results indicated that the marginal sinus and marginal zone were morphologically and functionally destroyed in the ΔSpE mouse spleen.

**Figure 5.**
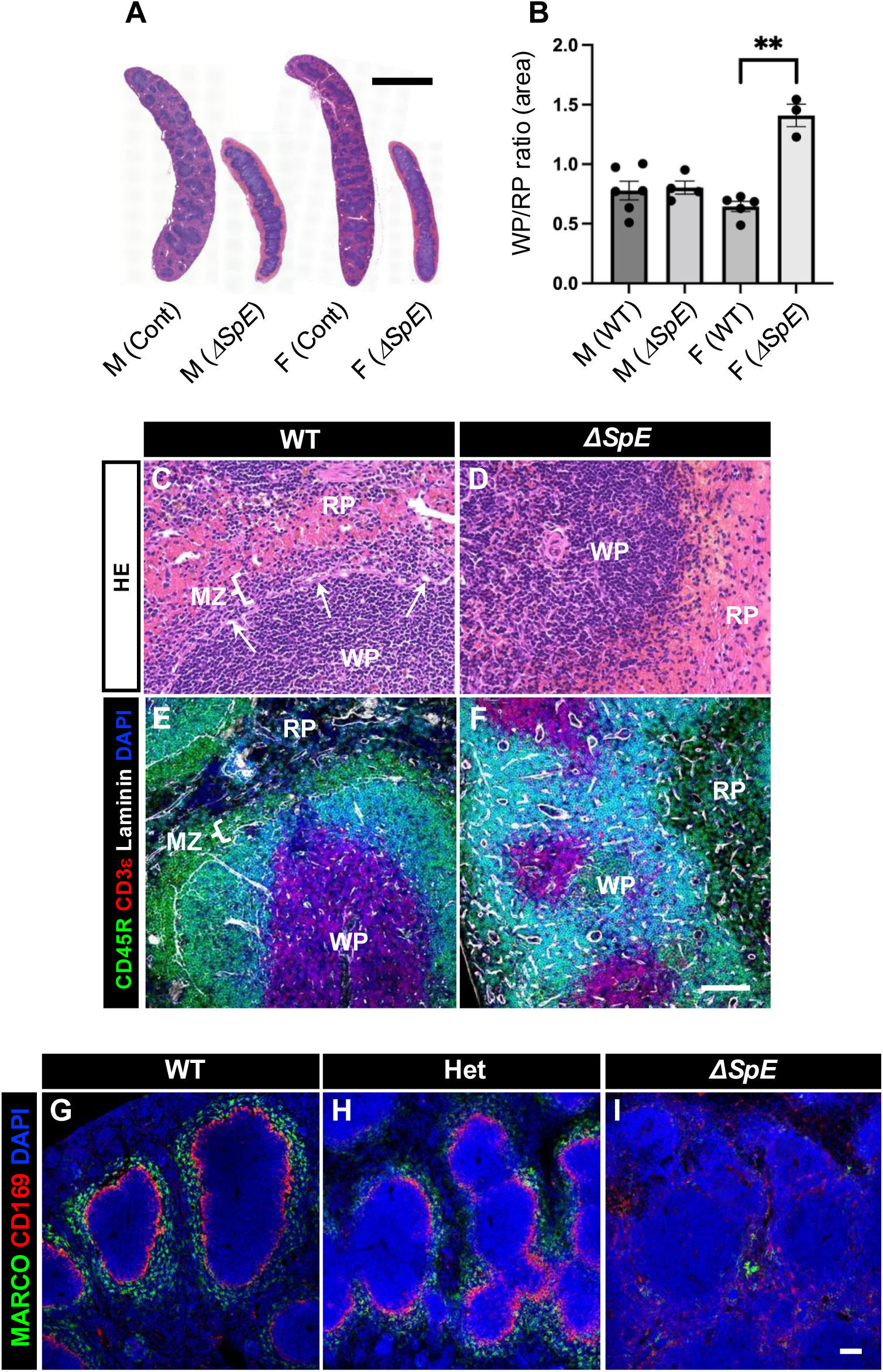
Morphological analysis of ΔSpE spleens. (A) Hematoxylin and eosin (HE) staining of spleens from each genotype and sex. Scale bar = 2 mm (B) White pulp-to-red pulp area ratios in each genotype and sex. Sample sizes: male wild-type, n = 6; male ΔSpE, n = 4; female wild-type n = 5; female ΔSpE, n = 3. *p* < 0.01 (**). (C, D) High-magnification HE-stained images of wild-type (WT) and ΔSpE spleens. Arrows indicate marginal sinuses. Scale bar = 100 µm. (E, F) Immunostaining of CD45R, CD3ε, and laminin in wild-type (WT) and ΔSpE spleens. RP, red pulp; WP, white pulp; MZ, marginal zone. Scale bar = 100 µm. (G., H., I) Immunostaining for MARCO and CD169 in wild-type (WT), heterozygous (Het), and homozygous (ΔSpE) spleens. Scale bar = 100 µm.

### Decreased steroidogenic activity in ΔSpE spleen

Bulk RNA sequence analyses were performed to comprehensively explore the functional changes in the spleens of ΔSpE mice. We collected spleens from wild-type and ΔSpE mice (three males and three females per genotype) and performed RNA sequencing. In addition, EGFP-positive cells were isolated from the spleens of Ad4BP-BAC-EGFP mice (three males and three females) using FACS to obtain bulk RNA-seq data. First, we extracted genes whose expression was decreased in the spleens of ΔSpE mice compared to wild-type mice. Notably, the number of downregulated genes in females (1,059) was approximately twice that in males (552). Given this pronounced sex difference, subsequent analyses were performed separately for males and females, and only genes consistently detected in both sexes were retained for further examination.

Next, we compared the transcriptomes of whole-spleen tissues from wild-type mice with those of FACS-isolated NR5A1-positive cells (<u>s</u>plenic <u>e</u>ndothelial <u>c</u>ells; SECs). We found that 3,481 genes in males and 3,765 genes in females were expressed at higher levels in SECs. Among these, genes that were both highly expressed in SECs and downregulated in ΔSpE spleens numbered 159 in males and 132 in females, with 88 genes shared between the sexes (Fig. 6A, 6B; gene list is shown in Supplemental Table 4). GO analysis of these 88 genes revealed enrichment for terms related to the regulation of adrenal cortical hormone production, suggesting a strong link between adrenal tissue and NR5A1 (Fig. S8). Indeed, the list included *Star* and *Cyp21a1*, both of which are essential for adrenal corticosteroid synthesis (Fig. 6C, 6D, 6E).

**Figure 6.**
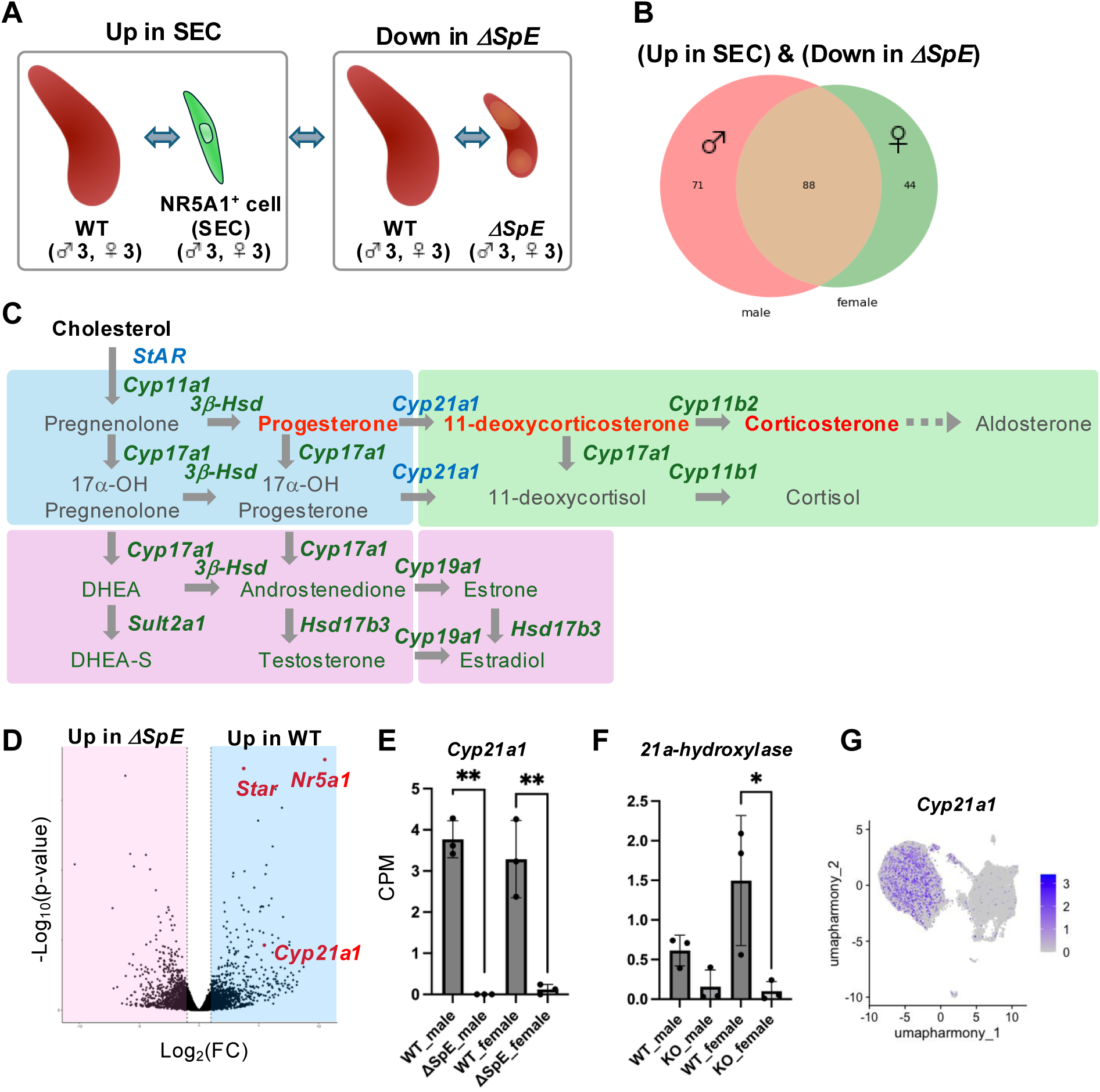
Downregulated genes in ΔSpE spleens. (A) Schematic representation of the strategy for gene expression analyses. Comparison of gene expression between wild-type (WT) spleens and sorted NR5A1^+^ sinusoidal endothelial cells (SECs) identified genes enriched in SECs (Up in SEC). Comparison between WT and ΔSpE spleens identified genes downregulated in ΔSpE (Down in ΔSpE). Genes included in both categories (Up in SEC and Down in ΔSpE) were extracted for each sex. (B) Venn diagram showing overlapping (Up in SEC and Down in ΔSpE) genes in both sexes. (C) Schematic representation of steroid biosynthetic pathways, including steroidogenic enzymes, precursors, and metabolites. Enzymes indicated in blue are among 88 genes identified in (B). Metabolites indicated in red represent corticosteroid hormones and their precursors. (D) Volcano plot analysis of gene expression in WT and ΔSpE spleens. Genes in red indicate representative genes downregulated in ΔSpE. (E) Expression levels of *Cyp21a1* gene in WT and ΔSpE spleens in each sex, showing as counts per million (CPM). *p* < 0.01 (**). (F) Metabolic activity of 21α-hydroxylase, assessed by precursor and metabolite concentrations in WT and ΔSpE spleens of each sex, determined by gas chromatography-mass spectrometry (GC-MS). *p* < 0.05 (*). (G) Feature plot of *Cyp21a1* expression in a UMAP presentation of NR5A1^+^ endothelial cells in the spleen.

We focused on CYP21A1, an enzyme essential for adrenocortical steroid hormone production and confirmed that *Cyp21a1* expression was almost completely lost in both male and female ΔSpE spleens (Fig. 6E). Splenic steroids were quantified using LC-MS, and the concentrations of 11-deoxycorticosterone and its precursor progesterone were determined (Fig. S9A). The metabolic ratio of these hormones was then calculated, which revealed a slight and considerable reduction in CYP21A1 enzymatic activity in males and females, respectively (Fig. 6F).

Single-cell transcriptomic analysis further demonstrated that *Star* and *Cyp21a1* were expressed in the endothelial cells of the splenic sinuses, whereas other genes required for adrenal corticosteroid biosynthesis (*Cyp11a1*, *Cyp17a1*, *Hsd3b1*, *Cyp11b1*, *Cyp11b2*) were minimally expressed or not expressed in NR5A1-positive cells (Fig. 6G, Fig. S9B).

Collectively, these findings indicate that splenic sinus endothelial cells possess StAR and CYP21A1 activities and contribute to *in situ* corticosteroid synthesis within the spleen. Moreover, the expression of steroidogenic genes in the spleen was dependent on NR5A1 expression.

### Lymph node-like structures and fibrosis in ΔSpE spleens

Compared with wild-type spleens, ΔSpE spleens showed the upregulation of 798 genes in males and 726 genes in females, with 427 overlapping genes between the sexes (Fig. 7A, 7B; the gene list is shown in Supplemental Table 5). GO analysis of these genes revealed multiple terms related to cell migration and lumen formation (Fig. S10). Notably, this gene list included the lymphatic marker *Lyve1*, as well as *Glycam1* and *Emcn*, which are expressed in HEVs (Fig. 7C, 7D). Double immunostaining for LYVE-1 and MECA79 revealed virtually no detectable expression of either marker in the wild-type spleens, whereas LYVE-1-positive lymphatic endothelial cells and MECA79-positive HEVs were clearly observed in the mesenteric lymph nodes. In the spleens of SpE heterozygous mice, MECA79 expression was detected mainly in the red pulp, whereas in ΔSpE mice, distinct LYVE-1-positive lymphatic-like structures and MECA79-positive HEV-like structures were evident (Fig. 7E–H). In contrast, double immunostaining for LYVE-1 and MAdCAM-1 showed that in wild-type spleens, MAdCAM-1 was expressed in the lining cells of the marginal sinus, whereas LYVE-1 expression was almost absent. However, in wild-type lymph nodes, both LYVE-1 and MAdCAM-1 were clearly expressed. In the spleens of SpE heterozygous mice, MAdCAM-1 expression was observed in the red pulp, and in ΔSpE mice, distinct LYVE-1-positive lymphatic-like and MAdCAM-1-positive HEV-like structures were apparent, showing a pattern similar to that in the mesenteric lymph nodes (Fig. 7I–L).

**Figure 7.**
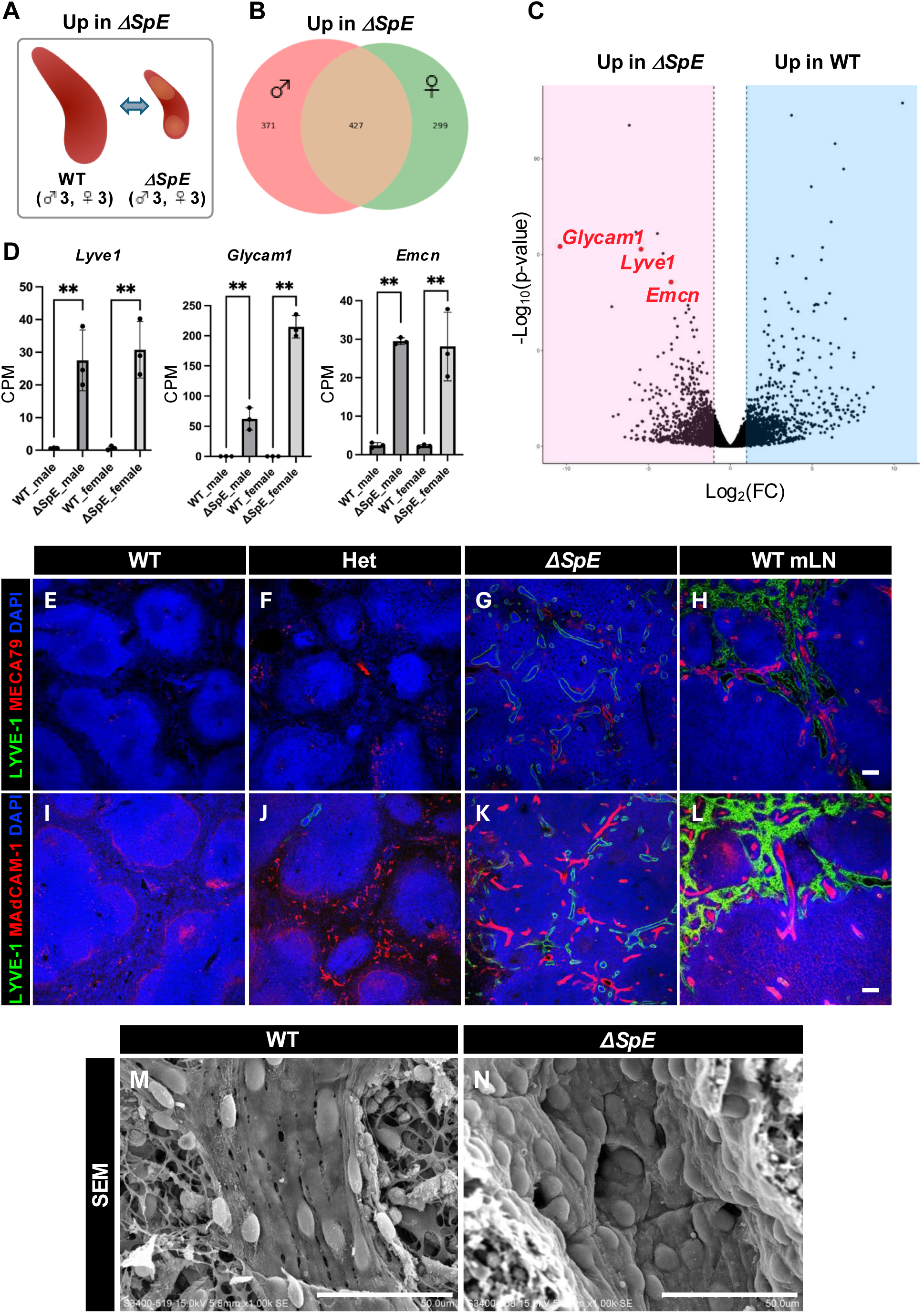
Upregulated genes in ΔSpE spleens. (A) Schematic representation of the strategy for gene expression analyses. Comparison of gene expression between wild-type (WT) and ΔSpE spleens identified genes upregulated in ΔSpE (Up in ΔSpE). (B) Venn diagram showing (Up in ΔSpE) genes in both sexes. (C) Volcano plot analysis of gene expression in WT and ΔSpE spleens. Genes in red indicate representative genes upregulated in ΔSpE. (D) Expression levels of *Lyve1*, *Glycam1*, *and Emcn* genes in WT and ΔSpE spleens in each sex, showing as counts per million (CPM). *p* < 0.01 (**). (E, F, G, H) Immunostaining of LYVE-1 and MECA79 in WT, heterozygous (Het), ΔSpE spleens and wild-type mesenteric lymph node (WT mLN). Scale bar = 100 µm. (I, J, K, L) Immunostaining of LYVE-1 and MAdCAM-1 in WT, heterozygous (Het), ΔSpE spleens and wild-type mesenteric lymph node (WT mLN). Scale bar = 100 µm. (M, N) Scanning electron microscopy (SEM) images of WT and ΔSpE spleens. Scale bar = 50 µm.

Scanning electron microscopy showed that in wild-type spleens, endothelial cells were elongated and rod-shaped, with characteristic intercellular slits, whereas in ΔSpE spleens, tall endothelial cells densely lined the vessel lumen, forming HEV-like structures without gaps (Fig. 7M, 7N). Additionally, the loss of fenestrated capillaries, a characteristic of the splenic sinuses, was associated with a significant increase in abnormal red blood cells in the peripheral blood, including Howell–Jolly bodies (Fig. S11). Collectively, these results indicate that ΔSpE spleens lose the identity of the spleen-specific vasculature and acquire a lymph node-like architecture.

We also found that the upregulated genes in ΔSpE spleens are related to “extracellular matrix (ECM) organization” and “elastic fiber formation” (Fig. S10). Double immunostaining for F4/80 and laminin revealed that in the spleens of SpE heterozygous mice, small vessels surrounded by laminin-positive basement membranes increased, whereas in ΔSpE mice, these small vessels developed at an even higher density, accompanied by a reduction in F4/80-positive macrophages in the red pulp (Fig. S12A–C). In addition, EVG staining suggested an increase in both collagen and elastic fibers in the spleens of ΔSpE mice. Furthermore, the lymphatic-like structures in the spleens of ΔSpE mice were surrounded by a thin collagen layer (Fig. S12D–I).

### NKX2.3 regulates Nr5a1 expression via direct binding to the SpE

Within the SpE, we identified several small subregions with sequences conserved across mammalian species (Fig. S5B). One of these, designated R2, contains a binding motif for the homeobox transcription factor NKX2.3. To test its function, we generated mice in which the R2 sequence was replaced with a mutated sequence containing a *Bam*HI restriction site using genome editing, which we named SpE-R2m (Fig. 8A, 8B). The spleens of SpE-R2m mice were markedly smaller than those of wild-type controls (Fig. 8C) and showed complete loss of NR5A1 expression (Fig. 8D–E). Moreover, the marginal sinus and marginal zone were absent (Fig. 8F–H), and ectopic lymphatic- and HEV-like structures were observed, which were essentially indistinguishable from those in ΔSpE mice (Fig. 8I–K). Notably, it has been reported that *Nkx2.3* knockout mice also display small spleens, loss of the marginal zone, and emergence of lymph node-like structures, closely resembling the phenotype of SpE-R2m mice^21,22,50^. In addition, our single-cell transcriptome data indicated that Nkx2.3 is expressed in both the splenic and marginal sinus endothelium (Fig. S13A), and its expression was unaffected by NR5A1 deletion in the spleen (Fig. S13B). Together, these findings strongly suggest that in splenic endothelial cells, NKX2.3 directly regulates *Nr5a1* expression through its binding to SpE.

**Figure 8.**
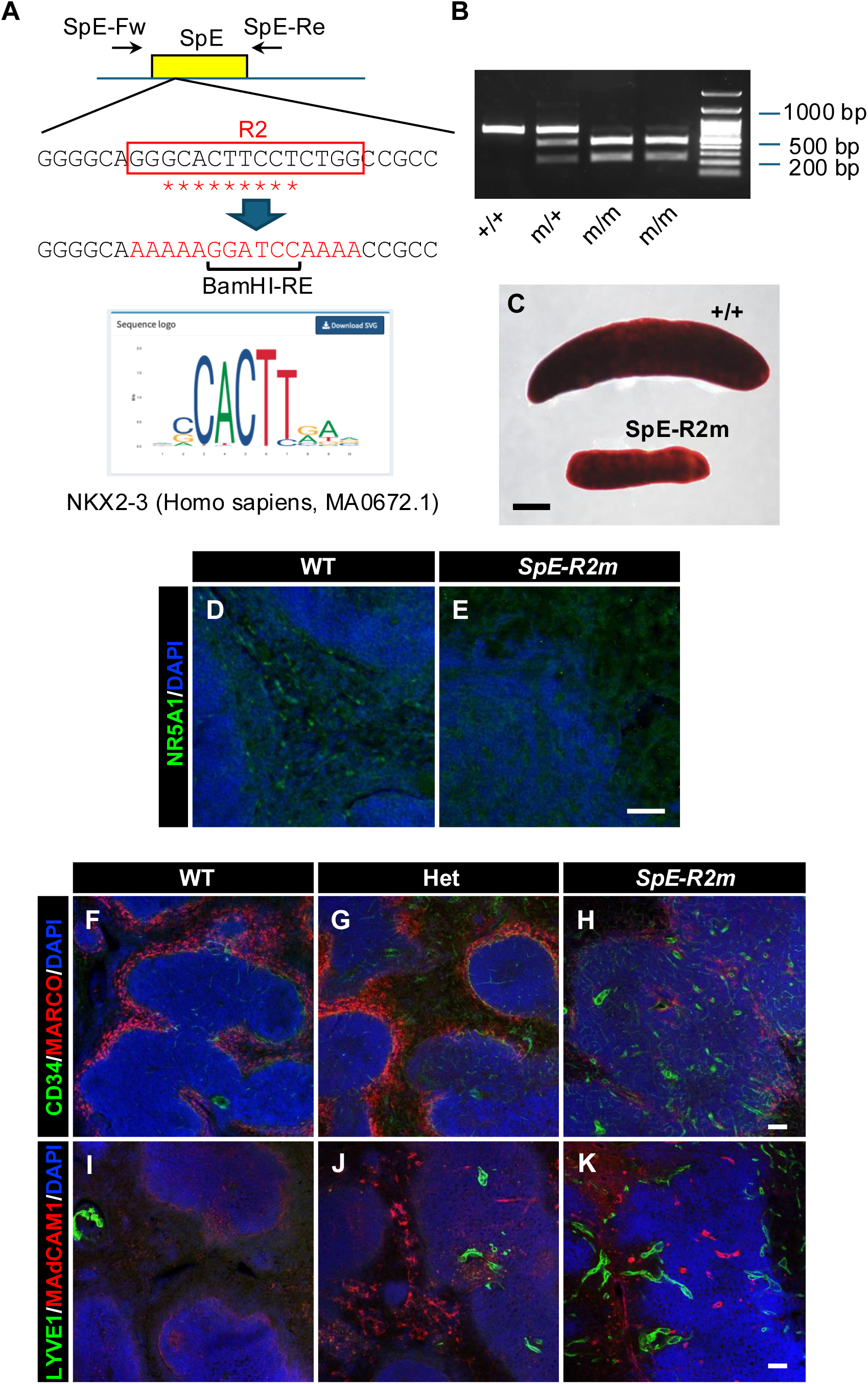
NKX2.3 regulates *Nr5a1* gene expression in the spleen via direct binding to SpE. (A) Schematic representation of a mutation induced in R2 sequence in SpE. The yellow box indicates SpE, and arrows indicate genotyping primers (SpE-Fw and SpE-Re). The red open rectangle indicates the R2 sequence, and asterisks indicate the NKX2.3-binding motif. Mutated R2 sequence (R2m) contains *Bam*HI restriction element (*Bam*HI-RE). The consensus sequence for human NKX2-3 binding was searched in JASPAR (https://jaspar.genereg.net/) and is shown in PWM format. (B) PCR-based genotyping of SpE-deleted mice. The wild-type allele yielded a 655-bp fragment, whereas the SpE-R2m allele yielded 429-bp and 225-bp fragments. (C) Gross morphology of spleens from wild-type (+/+) and homozygous mutation in SpE (SpE-R2m) mice. Scale bar = 2 mm. (D, E) Immunostaining of NR5A1 in WT and SpE-R2m spleens. Scale bar = 50 µm. (F, G, H) Immunostaining of CD34 and MARCO in WT, heterozygous (Het) and homozygous SpE-R2m spleens. Scale bar = 100 µm. (I, J, K) Immunostaining of CD34 and MARCO in WT, heterozygous (Het), and homozygous SpE-R2m spleens. Scale bar = 100 µm.

## Discussion

### Heterogeneity of NR5A1^+^ endothelial cells in the splenic sinus and marginal sinus

The spleen has two major physiological functions closely associated with the vascular system. First, it acts as a filter that captures foreign substances and senescent red blood cells from circulation. This filtering function is mediated by the unique architecture of the splenic sinus, in which elongated rod-shaped endothelial cells are aligned longitudinally to form large intercellular gaps^1^. The second function is immunity, wherein lymphocytes densely clustered in the white pulp play a central role. The marginal sinus is located at the boundary between the white pulp and marginal zone, and it has been proposed that blood-borne foreign antigens and bacteria are captured by antigen-presenting cells residing in the marginal zone, thereby initiating immune responses^3,8,9,11,51,52^.

Although NR5A1 expression in splenic endothelial cells has been reported previously^14^, this study is the first to demonstrate its expression in both splenic and marginal sinus endothelial cells. Considering the functional division of the splenic and marginal sinuses described above, it is reasonable to assume that their endothelial cell properties are distinct.

Single-cell transcriptomic analyses detected these two populations as separate clusters. Simultaneously, our single-cell analyses also provided novel insights into their functional differences: marginal sinus endothelial cells exhibited higher expression of genes associated with vascular patterning and ECM degradation and remodeling, whereas splenic sinus endothelial cells showed elevated expression of genes related to angiogenesis and cell–substrate adhesion. Moreover, the observation that *Cd34*, a marker highly expressed in immature endothelial cells or endothelial progenitors^53^, is more strongly expressed in marginal sinus endothelial cells suggests that they represent a more primitive population, whereas splenic sinus endothelial cells are more differentiated and specialized. The developmental relationship between these two endothelial populations remains an important subject for future investigation.

### Downstream genes of Nr5a1 in the spleen

Gene expression analysis of ΔSpE spleens revealed reduced expression of *Star* and *Cyp21a1*, key enzymes in adrenal steroid hormone synthesis. Given that NR5A1 directly regulates steroidogenic enzymes in adrenocortical cells, it is plausible that NR5A1 also directly controls *Star* and *Cyp21a1* expression in the splenic sinus endothelial cells.

Although StAR and CYP21A1 are expressed in NR5A1^+^ splenic cells, no other enzymes required for complete adrenal steroidogenesis were detected. Nevertheless, adrenal steroids were clearly present in whole-spleen extracts, implying that additional cell types contributed to the remaining enzymatic steps. To our knowledge, this is the first report of adrenal steroid hormone production in the spleen, although its physiological relevance remains to be elucidated.

While NR5A1 appears to be indispensable for steroidogenesis in the spleen, the marked disruption of the splenic vascular architecture in ΔSpE mice suggests that NR5A1 orchestrates the morphological and functional differentiation of splenic endothelial cells. Thus, NR5A1 may regulate distinct gene networks in the splenic and marginal sinuses. Our single-cell analyses indicated that *Star* and *Cyp21a1* are NR5A1 targets in the splenic sinus, whereas the downstream effectors of NR5A1 in the marginal sinus remain unclear.

Immune cells localized in the marginal zone are responsible for mounting immune responses against blood-borne antigens and play a crucial role in the immunological function of the spleen. Our findings suggest that defective formation of the marginal sinus disrupts the microenvironment essential for the development of marginal zone macrophages and marginal zone B cells. Recent studies have shown that macrophages in the marginal zone can migrate and promote cardiac recovery after myocardial infarction by suppressing inflammation and fibrosis^12^. Therefore, NR5A1 is likely to be essential for establishing the tissue organization of the marginal zone that supports these functions. Elucidating how NR5A1 regulates the development and maintenance of the marginal sinus may provide new insights into the maintenance of immune homeostasis and the pathogenesis of immune-related diseases.

### Regulation of Nr5a1 gene expression in spleen

Both the spleen and lymph nodes belong to the category of secondary lymphoid organs, and the gut-associated lymphoid tissue (GALT) exhibits a structure similar to that of the lymph nodes. NKX2.3 is expressed in all these lymphoid organs, and analyses of *Nkx2.3* knockout mice have demonstrated that it plays essential roles in each of them. In contrast, NR5A1 expression was restricted to the spleen and was not detected in the lymph nodes or GALT. In this study, we demonstrated that the binding of NKX2.3 to the SpE of the *Nr5a1* gene is indispensable for the expression of *Nr5a1* in the spleen. However, the precise mechanism underlying the spleen-specific activity of SpE remains unclear. It is possible that sequences within SpE other than the R2 region contribute to the spleen-specific regulation of *Nr5a1*. Conversely, although SpE is required for NR5A1 expression in the spleen, we cannot exclude the possibility that additional regulatory regions outside the SpE within the *Nr5a1* locus, known as shadow enhancers, may also be involved in splenic expression. Previous studies have shown that the Polycomb factor M33 participates in the regulation of NR5A1 expression in both the spleen and adrenal gland^54^, raising the possibility of a functional relationship between M33 and SpE, which warrants further investigation.

### Gene dosage of Nr5a1 and spleen development

Our results demonstrated that heterozygous deletion or mutation of SpE produces a milder phenotype than homozygous deletion, indicating that NR5A1 expression levels are quantitatively important for spleen development. This observation is consistent with that of a previous report and supports the concept that NR5A1 dosage can exert dose-dependent effects on cell differentiation and tissue morphogenesis^55^. However, the phenotype of SpE heterozygous deletion mice was much milder than that of homozygous deletion mice. Similar observations have been reported for other genes, in which the deletion of only one pair of enhancers resulted in a markedly attenuated phenotype compared to homozygous deletion. This phenomenon is generally understood to reflect redundant functions of enhancers^56^.

Clinically, NR5A1 haploinsufficiency in humans is associated with a spectrum of adrenal and gonadal phenotypes, including varying degrees of adrenal insufficiency and sex development^57,58^. Our data extend this concept to the spleen, demonstrating that subtle reductions in NR5A1 expression can also influence vascular and structural organization, and reinforcing the broader principle that the gene dosage of key transcription factors is a critical determinant of organ development and phenotypic variability.

Collectively, these results provide experimental support for the notion that transcription factor dosage is a general and quantitative mechanism that regulates cell differentiation and tissue morphogenesis, with NR5A1 representing a tissue-specific example of this mechanism in the spleen.

### Sex difference of steroidogenesis in the spleen

Even in wild-type mice, females have larger spleens and exhibit sex-specific differences in immune cell composition and circulating cytokine levels^49^. In this study, we further found that the effects of NR5A1 deficiency in the spleen differed between sexes, with a significantly increased white pulp-to-red pulp ratio in females. This is consistent with the marked reduction of CYP21A1 expression (and thus 21α-hydroxylase activity) in the red pulp of females.

Sex differences in immune function are well established in both mice and humans^59^. Because circulating adrenal steroid hormone levels also show sexual dimorphism, it is plausible that spleen-derived adrenal-type steroid production contributes to these sex differences in immune function and possibly to immune-related pathologies.

In our study, the SpE-deficient phenotype was more pronounced in female mice, suggesting that NR5A1-dependent splenic development is regulated in a sex-specific manner. One explanation may be that baseline NR5A1 expression or its developmental timing differs between sexes, such that the partial reduction in females drops more easily below the threshold required for proper endothelial-like cell differentiation and vascular organization. Alternatively, sex hormones, such as estrogen or androgens, may modulate NR5A1-dependent programs during development.

These findings are consistent with the broader concept that transcription factor dosage exerts sex-dependent effects on organogenesis. Similar dose- and sex-dependent phenotypes have been reported for other developmental regulators such as SOX9^60^ and GATA4/6^61^, where small differences in expression lead to divergent outcomes between the sexes. Thus, our results emphasize that both gene dosage and sex-specific context shape the robustness and variability of organ development.

From a translational perspective, these findings might be relevant to human NR5A1 haploinsufficiency, in which clinical manifestations, including adrenal and gonadal phenotypes, vary between the sexes. Our data suggest that subtle reductions in NR5A1 expression could similarly affect splenic development in a sex-dependent manner, thereby influencing immune function or disease susceptibility.

## Conclusions

In this study, we demonstrated that NR5A1 expression in splenic endothelial cells is governed by the SpE, which contains a functional NKX2.3-binding motif. Our findings reveal fundamental mechanisms underlying the establishment of organ-specific vascular identity and provide new insights into how transcription factor dosage and sex-specific regulatory programs shape tissue architecture. However, our study has some limitations. The developmental relationship between splenic and marginal sinus endothelial cell populations remains unclear, and the physiological significance of local steroid hormone production in the spleen requires further investigation.

The implications of our findings extend beyond basic developmental biology to potential clinical relevance. The disruption of marginal zone architecture in SpE-deficient mice impairs the microenvironment necessary for immune cell localization and function, suggesting that NR5A1 deficiency may influence immune homeostasis and susceptibility to immune-related diseases. The pronounced sex differences observed in SpE-deficient phenotypes, particularly in steroidogenic activity and red and white pulp organization, raise important questions about whether human NR5A1 haploinsufficiency similarly affects splenic development and immune function in a sex-dependent manner. Future studies should focus on identifying the complete spectrum of NR5A1 target genes in both splenic and marginal sinus endothelial cells, elucidating the functional consequences of spleen-derived steroid hormones on local and systemic immunity, and investigating whether NR5A1-dependent splenic development contributes to sex-specific differences in immune responses and disease susceptibility. Understanding these mechanisms may inform therapeutic strategies for immune disorders associated with splenic dysfunction.

## Acknowledgements

We appreciate the technical support provided by the Central Research Center, Kawasaki Medical School, and the Institute for Disease Modeling, Kurume University School of Medicine. We also thank the organizers of the Medical Research Center at Saitama Medical University for providing continuous support throughout this study. This study was supported by the Research Program for the Inter-University Research Network for High-Depth Omics, IMEG, Kumamoto University, Japan. This work was supported in part by the MEXT Promotion of Development of a Joint Usage/Research System Project: Pan-Omics DDRIC, MRCI for High-Depth Omics, CURE: JPMXP1323015486 for MIB, RIIT, and the Autonomous Medical Research Center at Kyushu University. We would like to thank Editage (www.editage.jp) for English language editing.

## Author contributions

Conceptualization, Y. S.; Investigation, K. M., K. O., Miki Inoue, T. S., A. Y., Masanori Iseki, Katsuhiko Ishihara, T. M., R. S., Kei-ichirou Ishiguro, J. N., M. H. C., T. B., Y. O., E. K., K. T., and Y. S.; Resources, T. I.; Format analysis and Writing – Original Draft Preparation, K. M., K.O., and Y. S.; Writing – Review & Editing, T. S., A. Y., Masanori Iseki, Katsuhiko Ishihara, T. M., Miki Inoue, R. S., Kei-ichirou Ishiguro, M. H. C., T. B., Y. O., E. K., K. T., and Y. S.; Funding Acquisition, K. M., K. O., Y. S., and Supervision, Y. S.

---

Supplemental Figure 1. **Comparison of endogenous NR5A1 and EGFP expression in the spleens of Ad4BP-BAC-EGFP mice.**

Double immunostaining for NR5A1 (red) and EGFP (green) and nuclear counterstaining with DAPI (blue) were performed on spleen sections from Ad4BP-BAC-EGFP mice. Images of the same field are as follows: (A) NR5A1 channel; (B) EGFP channel; (C) merged NR5A1 and EGFP; and (D) merged NR5A1, EGFP, and DAPI. Magnified views of the regions including the marginal sinus are presented as A’, B’, C’, and D’, with arrows indicating endothelial cells of the marginal sinus. Likewise, enlarged views of regions including the splenic sinus are shown as A’’, B’’, C’’, and D’’, with arrowheads indicating the endothelial cells of the splenic sinus. Scale bar = 50 µm.

Supplemental Figure 2 **Spatial relationship between EGFP-positive cells and MAdCAM-1-positive cells in the spleen of Ad4BP-BAC-EGFP mice.**

(A) Immunofluorescence image of MAdCAM-1, EGFP, and Laminin, together with DAPI nuclear staining, in the spleens of Ad4BP-BAC-EGFP mice. The region outlined by the yellow box is shown enlarged in panel B. Scale bar = 50 µm The image in panel B is further separated into individual channels: MAdCAM-1 (red) in panel C; laminin (white) in panel D; merged MAdCAM-1 and EGFP in panel E; EGFP (green) in panel F; and DAPI (blue) in panel G. Dotted lines in panels C, D, F, and G indicate MAdCAM-1-positive cells. (H) Line plot generated along the direction indicated by the arrow in Panel B. The plot shows that EGFP-positive cells (green) are adjacent to MAdCAM-1-positive cells (red). Scale bar = 10 µm

Supplemental Figure 3. **Gene expression profiles in each cluster of NR5A1+ cells in the spleens of Ad4BP-BAC-EGFP mice.**

(A) UMAP plot of cells from three mice (two males and one female) after merging the datasets, with cells colored according to their individual origin.

(B) Feature plots of representative genes commonly expressed in clusters 1 and 2.

(C) Feature plots of representative genes that were highly expressed in cluster 1.

(D) Feature plots of representative genes highly expressed in cluster 2.

(E) Feature plots of representative genes highly expressed in cluster 6 (highlighted with red circles).

Supplemental Figure 4. **GO analysis of genes enriched in each cluster of NR5A1^+^ sinus endothelial cells in the spleens of Ad4BP-BAC-EGFP mice.**

(A) Bubble plot showing the molecular functions associated with each cluster identified by scRNA-seq in EGFP^+^ sinus endothelial cells. (B) Bubble plot showing the cellular components associated with each cluster identified by scRNA-seq of EGFP^+^ sinus endothelial cells. Gene Ontology (GO) analysis was performed using the top 150 highly expressed genes in each cluster.

Supplemental Figure 5. **Schematic illustration of the SpE deletion by CRISPR/Cas9.**

(A) Two guide RNAs were designed to target the upstream and downstream regions of the SpE. Arrows (SpE-Fw and SpE-Re) indicate the positions of the genotyping PCR primers. Nucleotide sequences surrounding the guide RNA target sites are shown (WT, wild type; de, SpE deletion). Guide RNA target sequences are shown in red, and PAM sequences are underlined.

(B) Alignment of the nucleotide sequences of SpE regions in mice, rats, rabbits, pigs, and dogs using MAFFT (https://mafft.cbrc.jp/alignment/software/). Asterisks indicate nucleotides conserved among all species. Guide RNA target sequences are shown in red, and PAM sequences are underlined. Nucleotides boxed in red indicate the R2 sequence, the mutation of which abolished enhancer activity.

Supplemental Figure 6. **Morphology of NR5A1-expressing organs other than the spleen in ΔSpE mice.**

(A) Gross morphology of the adrenal glands in wild-type (WT) and ΔSpE mice.

(B) Gross morphology of the ovaries in wild-type (WT) and ΔSpE mice.

(C) Gross morphology of the testes in wild-type (WT) and ΔSpE mice.

Scale bar = 2 mm.

Supplemental Figure 7. **Flow cytometric analysis of splenic immune cells in ΔSpE mice.**

(A) Representative flow cytometry plot of the spleen. B: B cells (CD19^+^), MZB: marginal zone B cells (CD19^+^CD21/35^high^CD23^low/-^), FoB: follicular B cells (CD19^+^CD21/35^high^CD23^high^), T1B: Transitional B cells of type1 (CD19^+^CD21/35^-^CD23^-^), T2B: transitional B cells of type 2 (CD19^+^CD23^+^CD21/35^+^IgM^+^).

(B) Flow cytometry of B-cell subsets isolated from the spleen of a wild-type mouse. CD21/35^high^CD23^low/-^ (marginal zone B cells) cells constitute 6.91% of CD19^+^ B cells.

(C) Flow cytometry of B-cell subsets isolated from the spleen of an SpE heterozygous mouse. CD21/35^high^CD23^low/-^ cells constitute 7.38% of CD19^+^B cells.

(D) Flow cytometry analysis of B-cell subsets isolated from the spleen of a ΔSpE mouse. CD21/35^high^CD23^low/-^ cells constituted 0.66% of CD19^+^B cells.

(E) Frequencies of B-cell subsets in the spleen: B: B cells, FoB: follicular B cells, T1B: transitional B cells of type 1, T2B: transitional B cells of type 2, MZB: marginal zone B cells, WT: wild-type: Het, heterozygous deletion of SpE, ΔSpE: deletion of SpE,*: *p* < 0.05, ns: not significant.

Supplemental Figure 8. **GO analysis of genes downregulated in the spleens of ΔSpE mice.**

(A) Biological processes associated with the downregulated genes in ΔSpE spleens.

(B) Tissue specificity of genes downregulated in ΔSpE spleens.

(C) Transcription factors predicted to regulate the genes downregulated in ΔSpE spleens.

Supplemental Figure 9. **Steroid levels and expression of steroidogenic genes in the spleen.**

(A) The concentrations of progesterone and 11-deoxycorticosterone were measured by LC-MS. M, male; F, female; WT, wild type.

(B) Feature plots showing the expression of the genes required for adrenal steroid production. *Star* and *Cyp21a1* were expressed in cluster 1, whereas the expression of the other genes was very low or undetectable (ND).

Supplemental Figure 10. **Upregulated genes the spleens of ΔSpE mice.**

Biological processes identified by GO analysis of genes upregulated in the spleen of ΔSpE mice.

Supplemental Figure 11. **Abnormal peripheral erythrocytes in ΔSpE mice.**

(A) Giemsa-stained peripheral blood samples from control male mouse.

(B) Giemsa-stained peripheral blood samples from a male ΔSpE mice. Arrows indicate abnormal erythrocytes containing Howell–Jolly bodies. Scale bar = 20 µm.

(C) The proportion of erythrocytes containing Howell–Jolly bodies in control and ΔSpE mice of each sex. At least 1,000 erythrocytes were examined per specimen (n = 4). *p* < 0.01 (**).

Supplemental Figure 12. **Increased ECM expression in the spleens of ΔSpE mice.**

(A–C) Double immunostaining for F4/80 (red) and Laminin (green) with nuclear counterstaining using DAPI (blue) in the spleen sections. WT, wild type; Het, SpE heterozygous. Scale bar = 100 µm.

(D–F) Hematoxylin and eosin (HE) staining of spleens from WT and ΔSpE mice. Arrowheads in (E) and (F) indicate ectopic lymphatic-like structures filled with lymphocytes.

(G–I) Elastica van Gieson (EVG) staining of spleens from WT and ΔSpE mice. The arrowheads in (H) and (I) indicate ectopic lymphatic-like structures filled with lymphocytes.

Scale bar = 50 µm.

Supplemental Figure 13. ***Nkx2-3* expression in the spleens of ΔSpE mice.**

(A) Feature plot showing *Nkx2-3* expression in NR5A1^+^ cells isolated from the spleens of Ad4BP-BAC-EGFP mice. *Nkx2-3* is expressed in clusters 1 and 2.

(B) *Nkx2-3* expression in the spleens of wild-type (WT) and ΔSpE mice of both sexes, as revealed by mRNA-sequence analysis. Expression levels are expressed as counts per million (CPM).

