## Supplemental Figures for "The *Nkx2.3*–*Nr5a1* gene cascade plays a crucial role in spleen-specific vascular architecture and marginal zone formation"

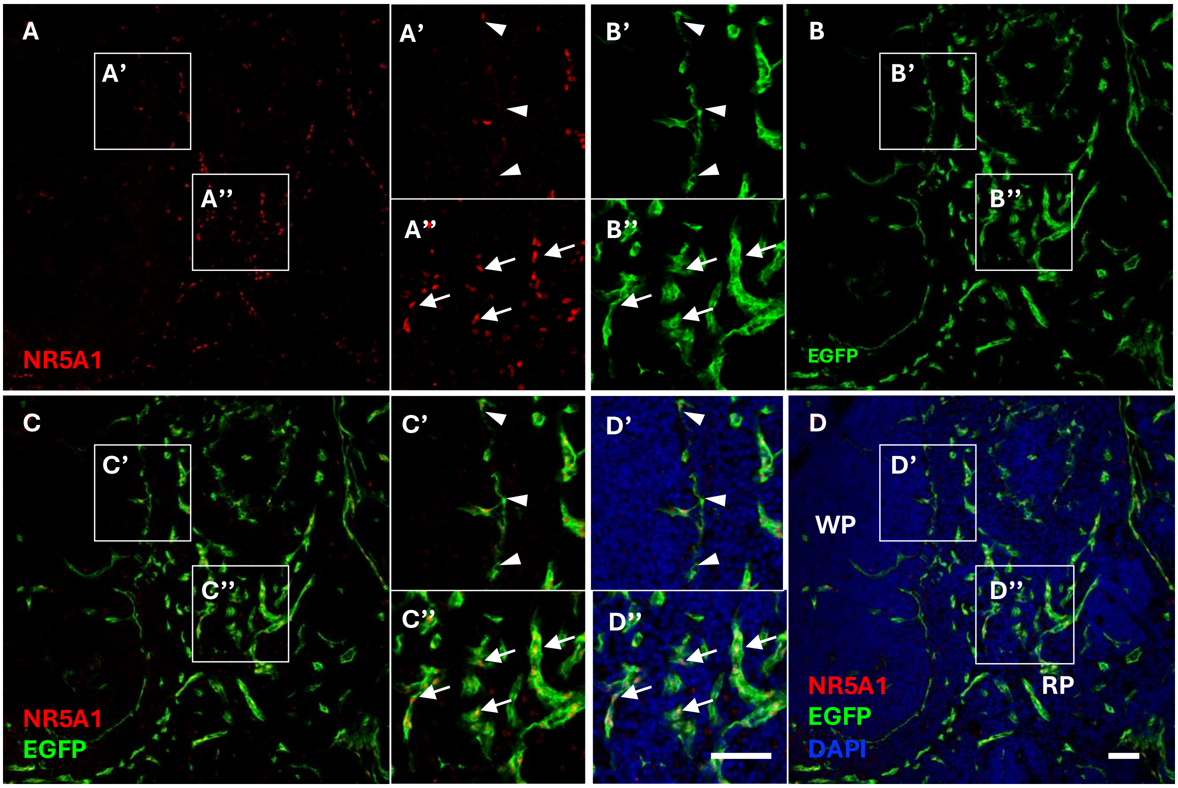


Miyabayashi, et al., Figure S1

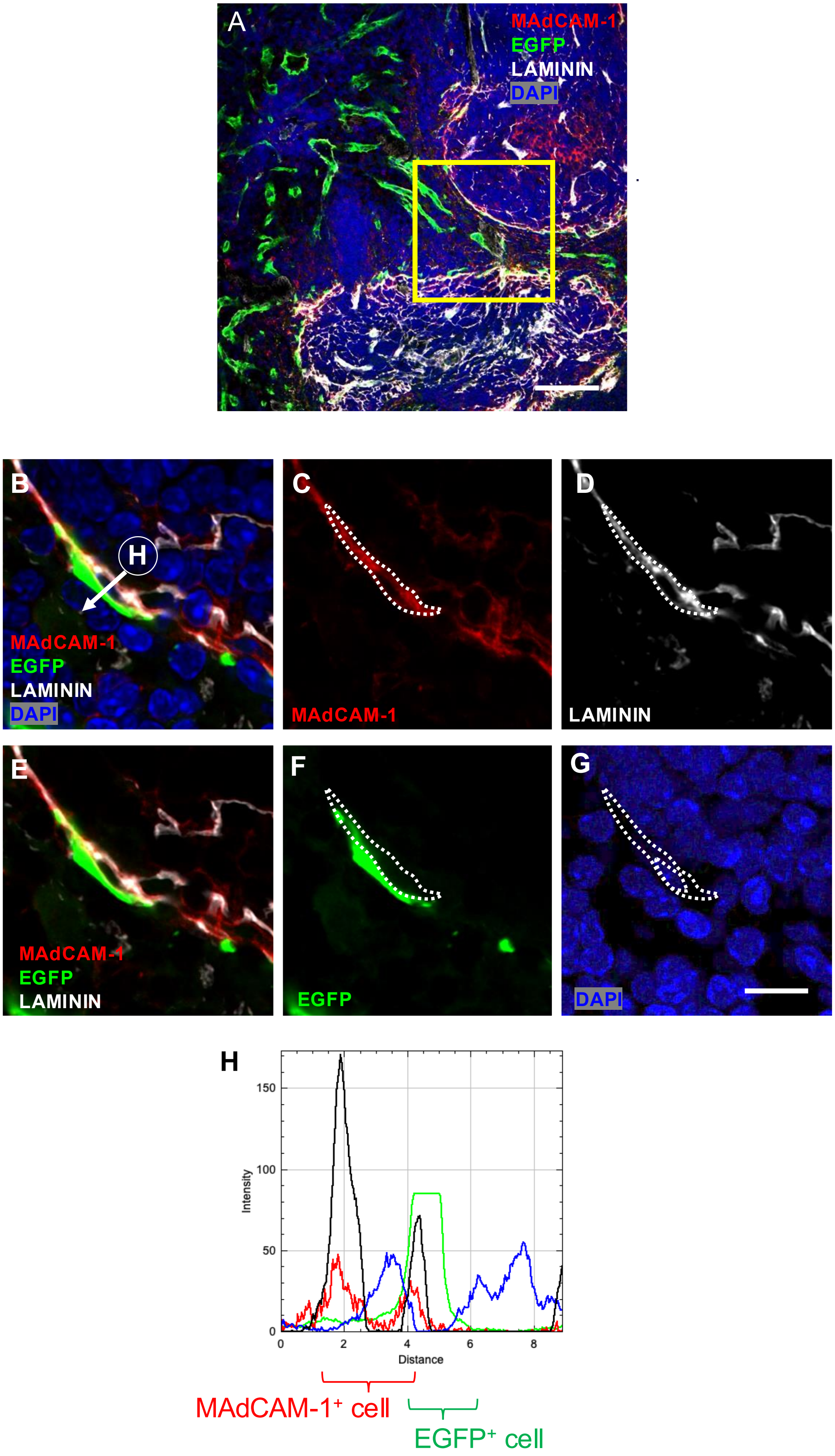


Miyabayashi, et al., Figure S2

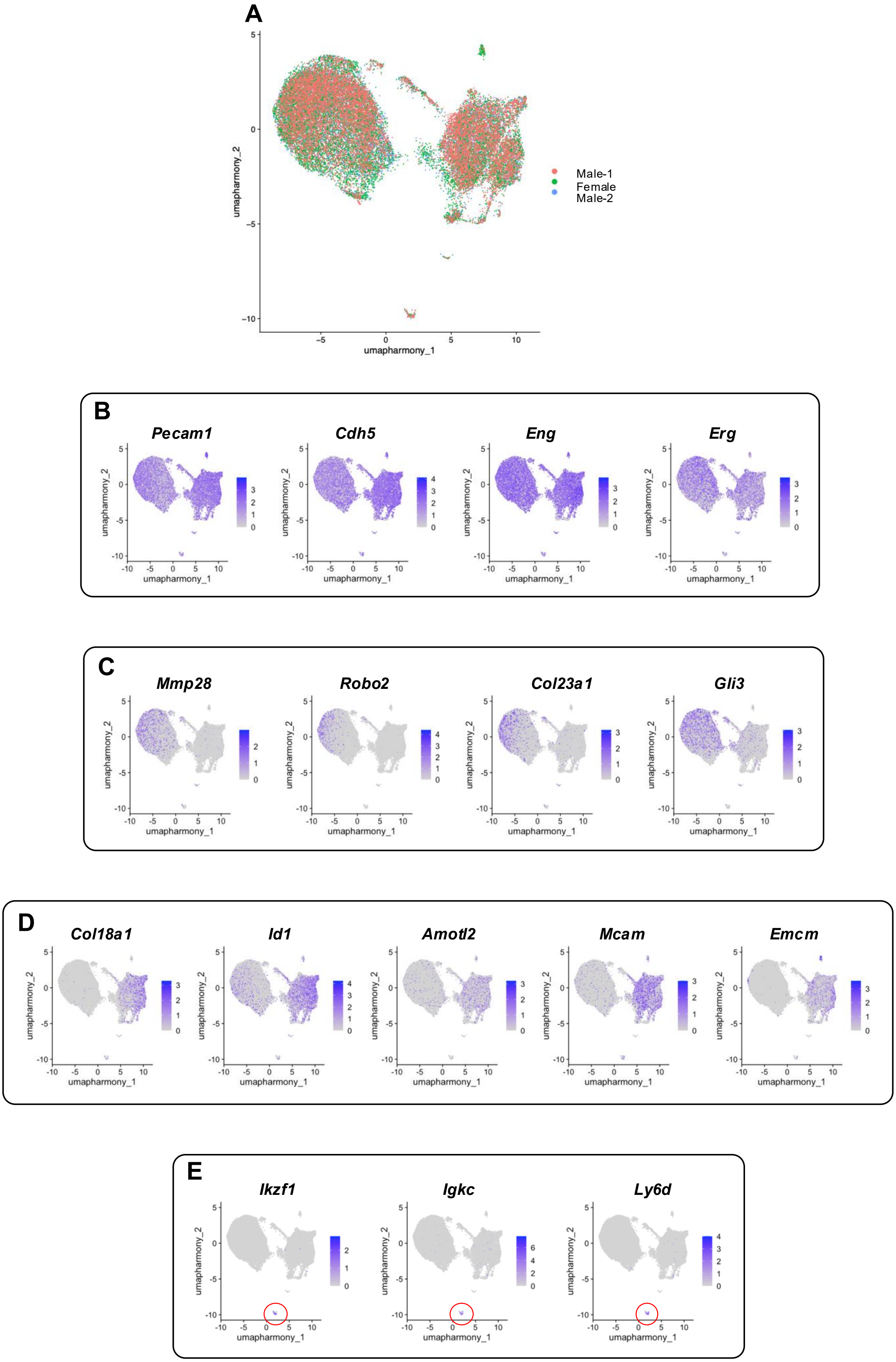


Miyabayashi, et al., Figure S3

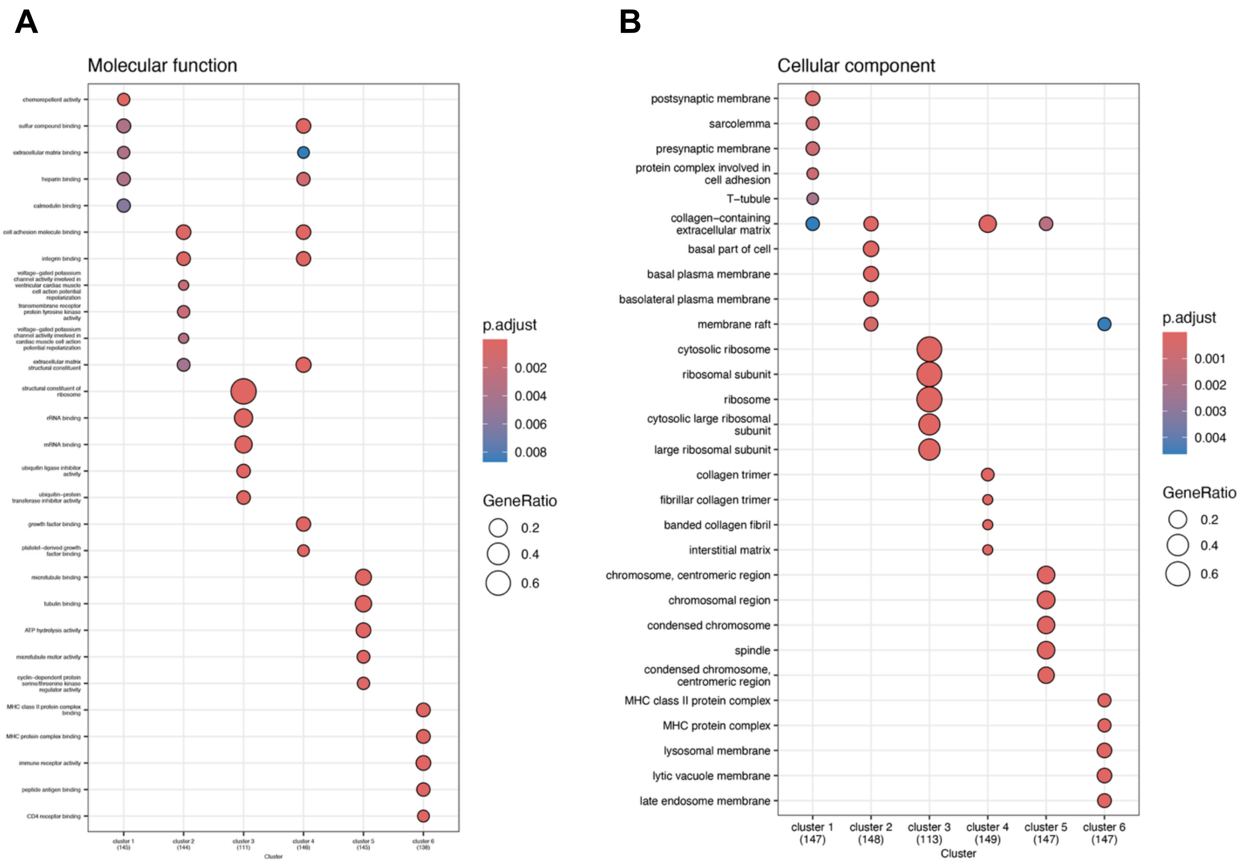


Miyabayashi, et al., Figure S4

**A**

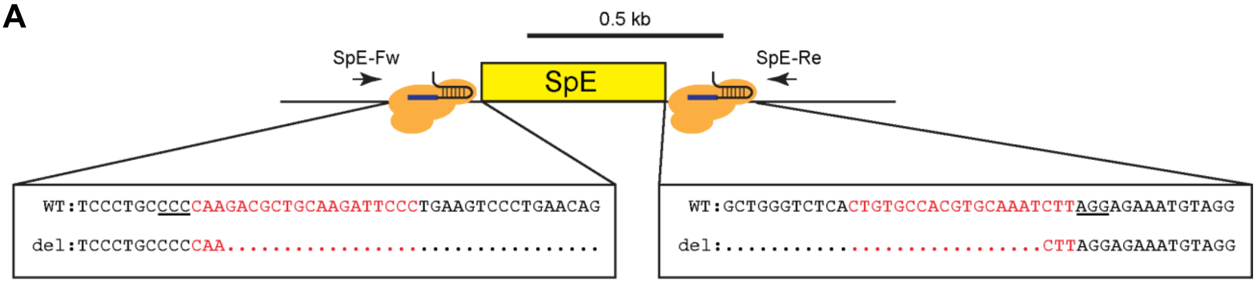


**B**

```
CLUSTAL format alignment by MAFFT (v7.490)

mouse_SpE       ---------tgctctggaagattctcctccctgccccaagacgctgcaagattccctga
rat_SpE         ggtgctcagtacgctggaagataccccagcccccacctcaagactgcaaacatcccccga
rabbit_SpE      ggagcttaag------------------------tcc-------ttcagacttcccatga
pig_SpE         ggatcctaagaccctggggttggttgccagcttcccc-------tccaggcttcccatga
dog_SpE         ggaccccaggacccttgggttggttgccagcttcccc-------tccagacttcccatga
                                                   **       .   ..  *.** .**

mouse_SpE       agtccc-tgaacagagagtagaaggagctaagcattctcagctttcctggatcttggatt
rat_SpE         agtccc-tgaacagagagtagaagaagctaag--------------------cttggatc
rabbit_SpE      gccccc---tgtgcagaggggaggagcagaacattcccagccttccagaaccctggatc
pig_SpE         gccccc------agtgcaggggaggagcagagcattcctggccttccagaaccctggatg
dog_SpE         gcccccctgtgttgggggaggaggagcagagcattcctggctttccagaaccctggatc
                . .***         * . .*.**.*** .*.                    *.*****

mouse_SpE       caggggcagggcacttcctctggccgccttcctggttggggaggtgggggtcatgcagg
rat_SpE         caggggcagggcacttcctctggctgccttcctggttggggaggtgggggtcacgcagg
rabbit_SpE      caggggcggggcacttcctctggccgccttcctgctctc-cgg--gggcttgcacgcagg
pig_SpE         caggggcagggcacttcctctggacgccttcctggtctctcgg--gggttttcacgcagg
dog_SpE         caggggcagggcacttcctctggccgccttcctagtctcttgg--ggttttttcacgcagg
                *******.*************** .********. *.    .*  **     **.*****

mouse_SpE       atgggaagggcttttgtcctagcgttgctgagtgtttttttcctgcttctggcttctctgt
rat_SpE         atgggaagggcttttgtcctagcgttgctgagtgtttttttcctgcttctggcttctctgt
rabbit_SpE      atgggaggggcttttgtcccagctccgctgagtgtttttttcctgcttctggcttctctgt
pig_SpE         atgggaagggcttctgtcccagcttcactgagtgtttttttcctgcttctggcttctctgt
dog_SpE         atgggaagggcttttgtcccagctttgctgagtgtttttttcctgcttctggcttctctgt
                ******.******.*****.*** ...*********************************

mouse_SpE       ggctcaaaccttcctgcttccccttcccagaacctgtgaaaggagcccaggaatcgaatc
rat_SpE         ggcccaaaccttcctgcttccccttcccagaacctgtgaaaggagcccagga-----atc
rabbit_SpE      ggtccaaaccttcctgcttcccctccccagagcctctgagcagagcccagg--------a
pig_SpE         ggtccaaaccttcctgcttccacttcccagagtctctgattcaagctcagga-----ata
dog_SpE         ggtccaaaccttcctgcttccccttcccagaacctctgagtcgagcccagga-----ata
                **..***************** **.******..** ***   .***.****

mouse_SpE       ttcaccata-gtcag-tgctgcttctgagggaccttgcagataagcccgagcccggcact
rat_SpE         ttagccata-ctcagttgctgcttgtgagggaccttgcagataagcccgagcccagtcct
rabbit_SpE      tgcaccctgccacccccacgttggtaaaggacccttgcagacaagcaccagcccagcacc
pig_SpE         tgaacca---ggtactaccctctacaaagaagccttgcaaataagtaccagcccagccct
dog_SpE         tgaaccatg-gttaccattctctgtaaaggagccttgcagataagtaccagcccagcctt
                *  .**       .    .  .    .**.. *******.*.***. * *****.*. ..

mouse_SpE       taca-gcctcgctagagccagcatgc-----------tttgccaggctgggggt-----
rat_SpE         tacg-gcct--cttgagcaggcatgc-----------tttgccaggct-ggggt-----
rabbit_SpE      tcga-acctc-acctgttggccatgtggg-------------------gaggggt---gc
pig_SpE         tcga-agcct-tctcatcagccatgtggaccttctttcccactggtctagcaagc---cc
dog_SpE         tagagggcct-cctaatgagccatgtggaacttctttcttgttggtctaacaagcaagcc
                *  . . *.   .  .   . ****.                       . ..*.

mouse_SpE       ---------ctgtgtcacagaggccaggcatgcaggtcagaagagctgaggctgcccagt
rat_SpE         ---------ctgtgtctaggaggccaggcatgcaggttggaagagctgaggttgcccagt
rabbit_SpE      tgggctgg-atctgattgcaaagttcg------ggacgaggcggtagaaaaccggggaca
pig_SpE         tgggctgatatctgtttgcaaagccag------agatagagaagcaggagaatgtgaggg
dog_SpE         ctggctagtgtctgtttgcaaagccag------aagtgggaaagcaggaaagtgggagta
                          * ** .   .*.*.. *      .... ... ..   .*.. .*   .

mouse_SpE       gagctgggt--------------------ctcactgtgccacgtgcaaatcttaggagaa
rat_SpE         gagctggtt--------------------cccaccatgccacatgcaaatcttaggcgat
rabbit_SpE      ggactgagtggccctgttgcccctctgggctggccgtccagcgccccacctccgggtgga
pig_SpE         gagctgagtgactctgttttctctttcgactgaaccttcactcccttaccttcgggcag-
dog_SpE         gtgctgagtgaccctcttttcccttttgg-tagccattcacccccctacttcca------
                * .***. *                     . . . * *  . . . * .....
```

Miyabayashi, et al., Figure S5

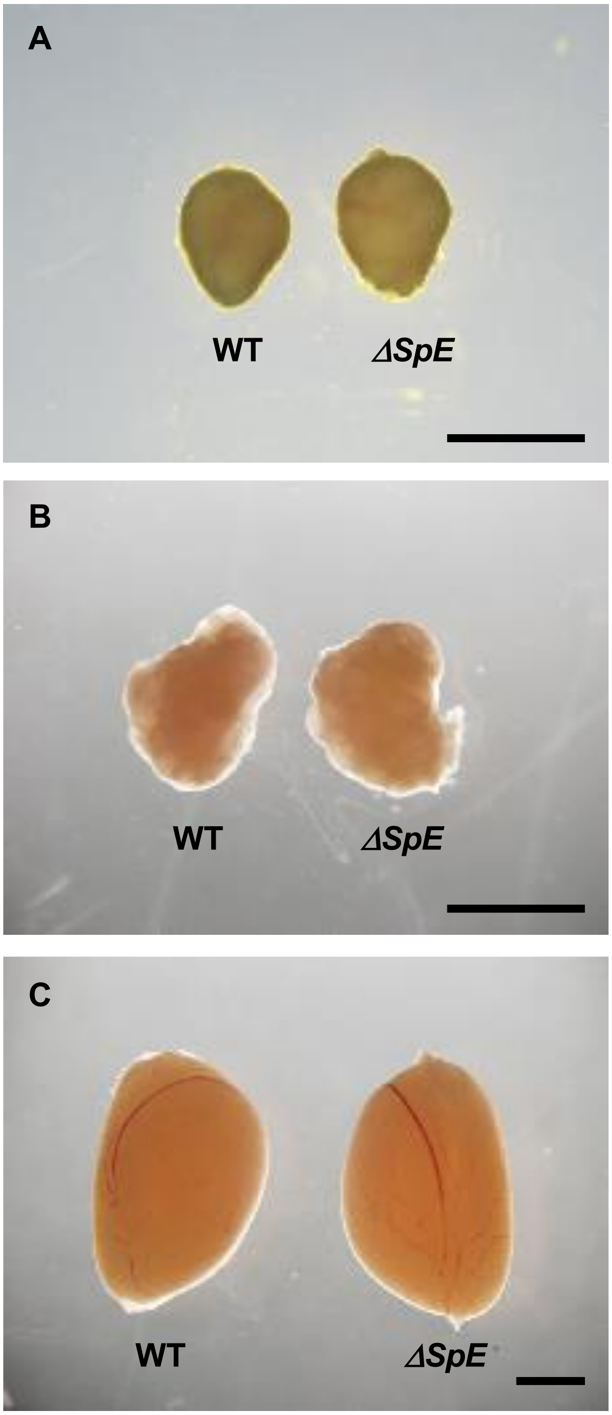


Miyabayashi, et al., Figure S6

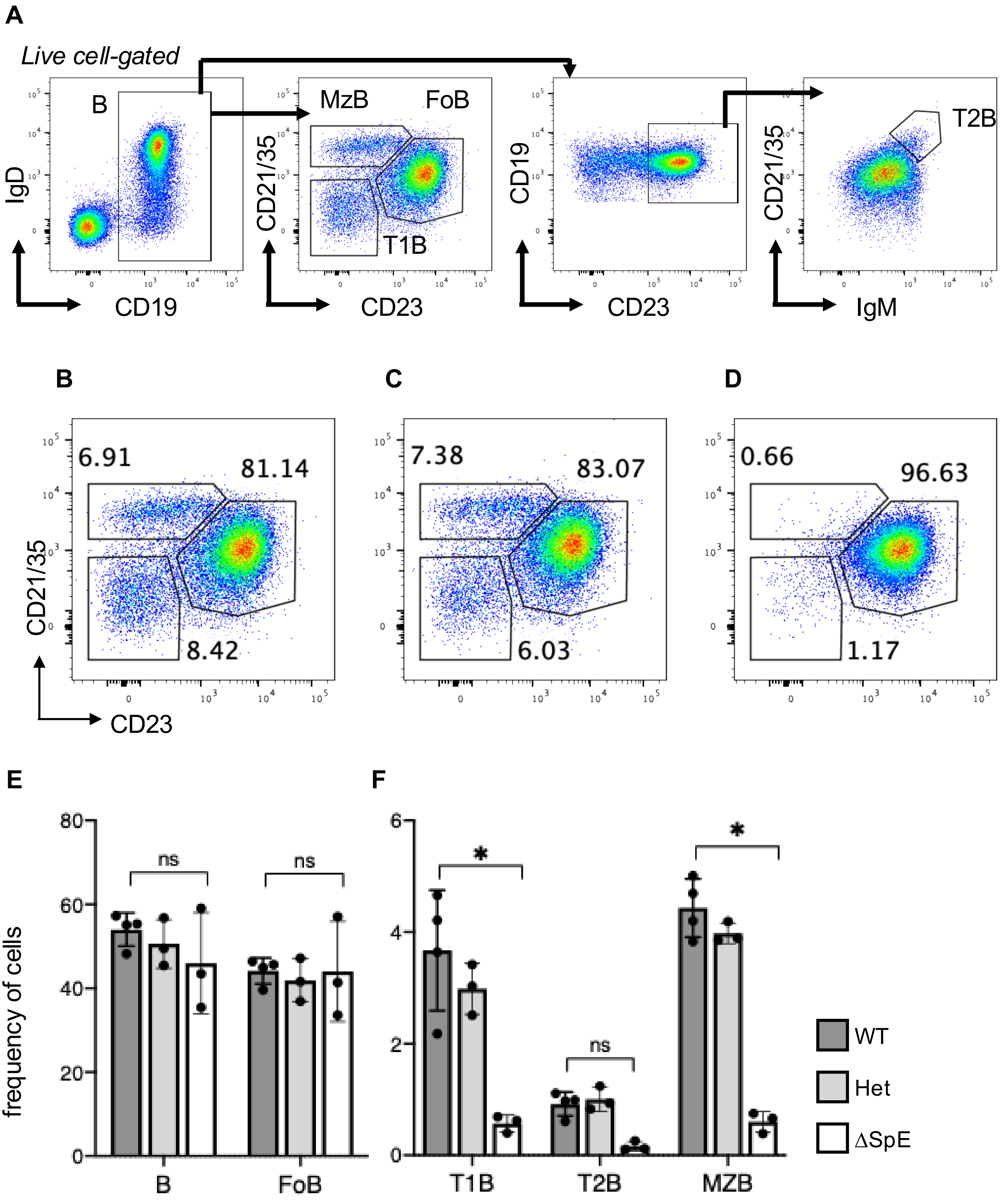


Miyabayashi, et al., Figure S7

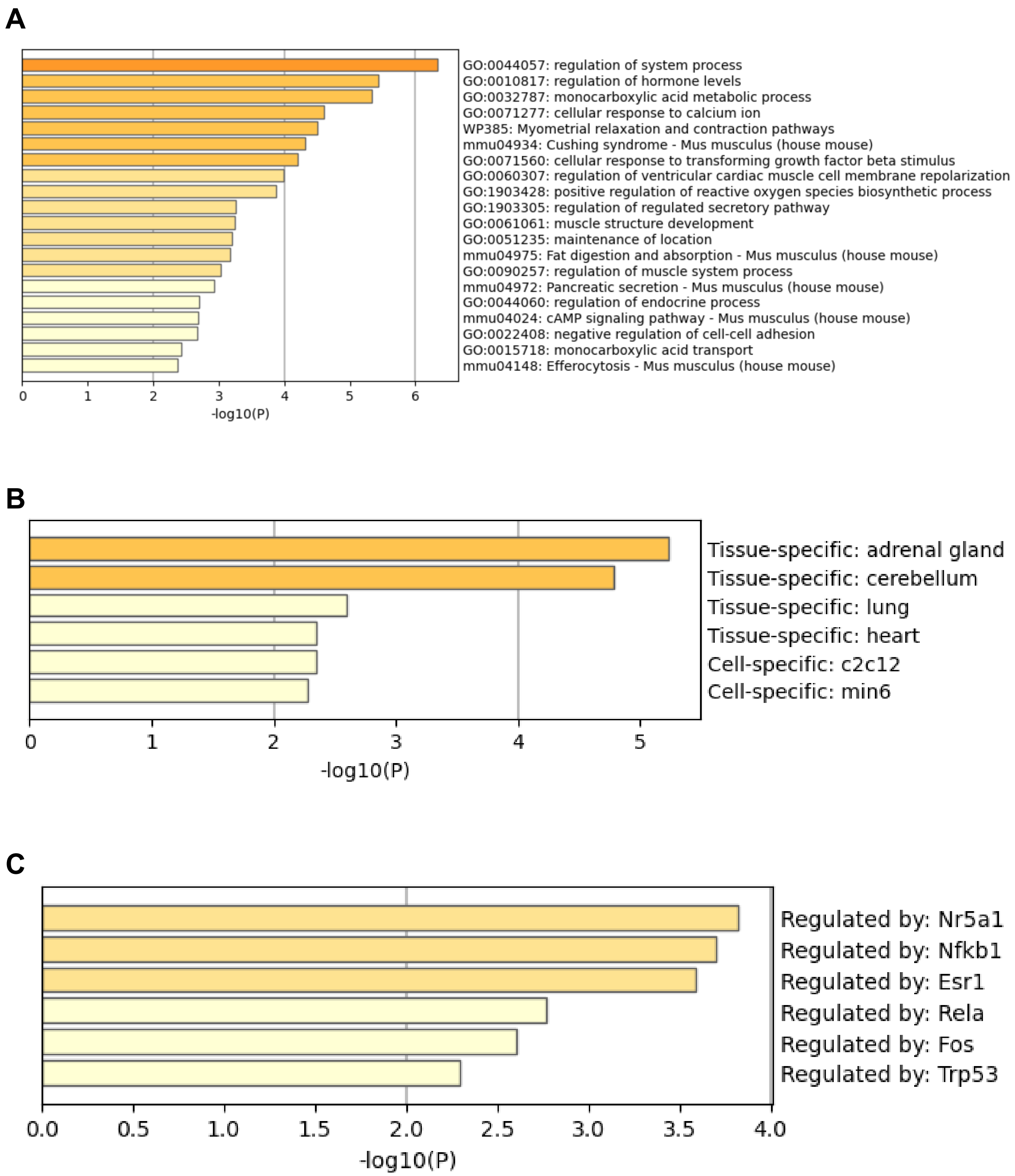


Miyabayashi, et al., Figure S8

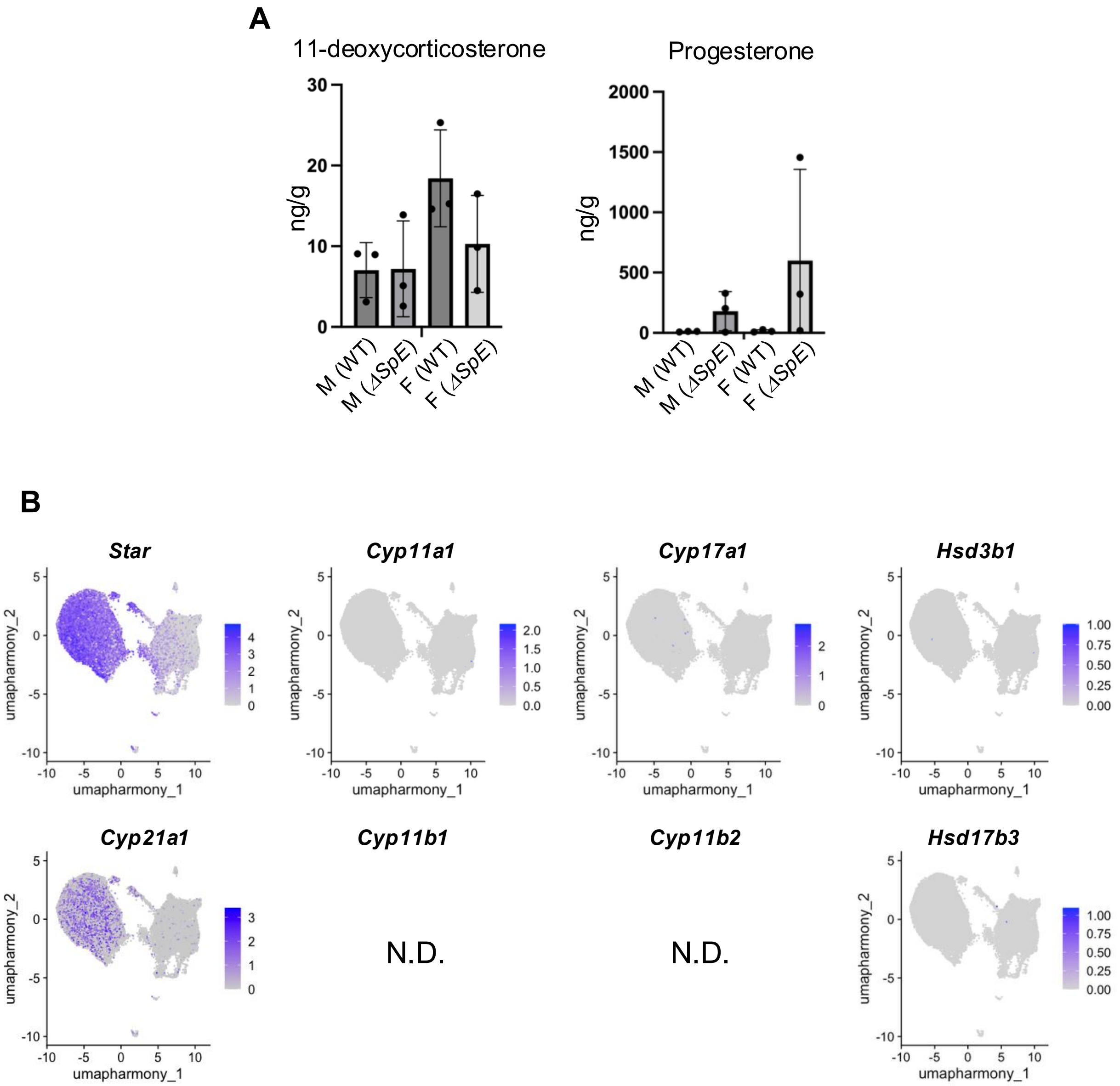


Miyabayashi, et al., Figure S9

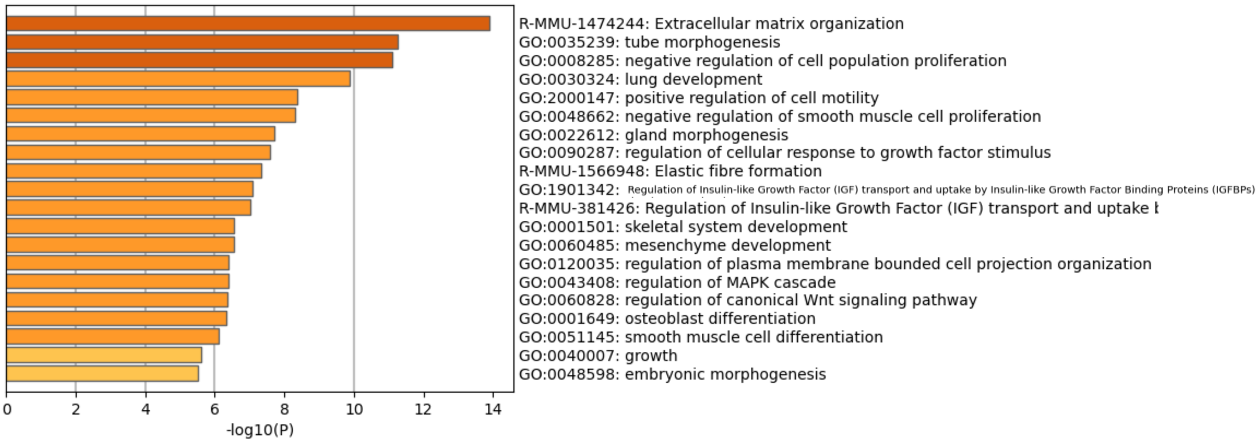


Miyabayashi, et al., Figure S10

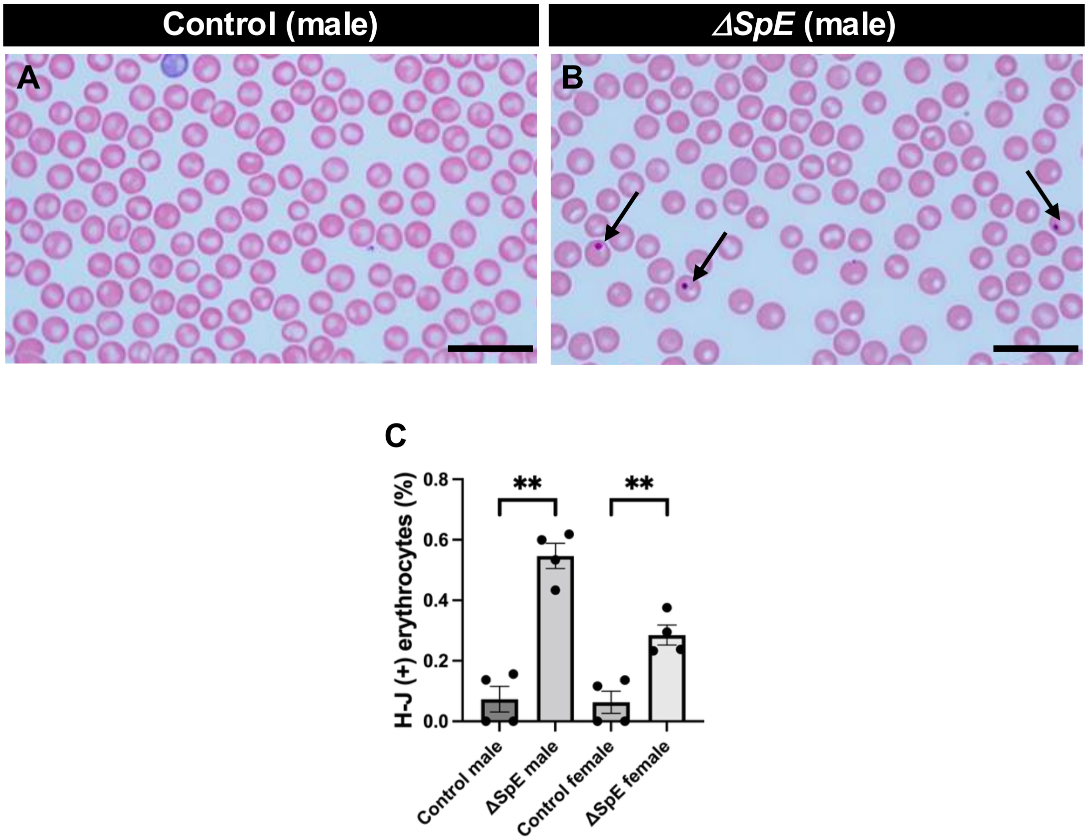


Miyabayashi, et al., Figure S11

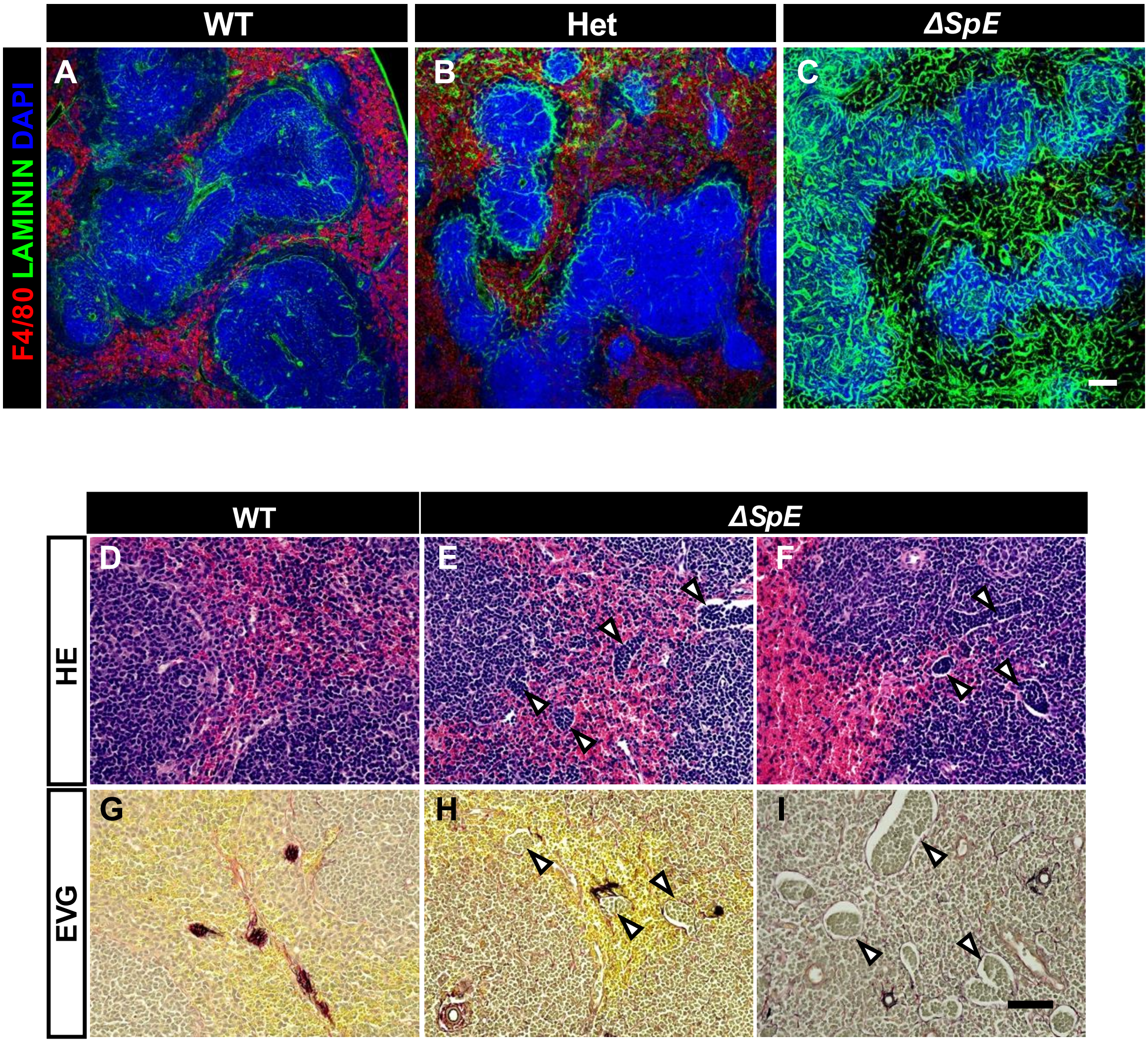


Miyabayashi, et al., Figure S12

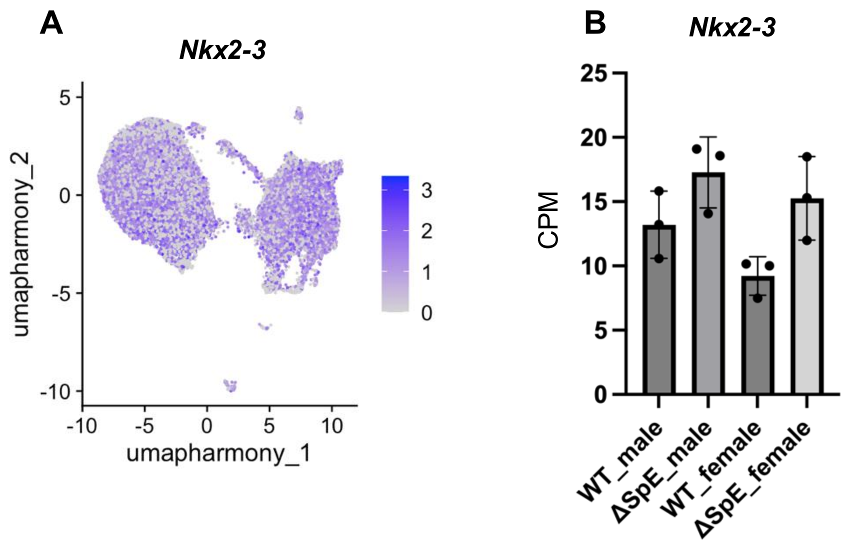


Miyabayashi, et al., Figure S13
