## Supplemental Table 1 for "The *Nkx2.3*–*Nr5a1* gene cascade plays a crucial role in spleen-specific vascular architecture and marginal zone formation"

Target sequence of guide RNAs

| Guide RNA | Target sequence |
|---|---|
| SpE-gRNA-up | GGGAATCTTGCAGCGTCTTGggg |
| SpE-gRNA-down | CTGTGCCACGTGCAAATCTTagg |
| SpE-R2mut | CCAACCAGGAAGGCGGCCAGagg |

Sequence of donor DNA for R2 mutagenesis

| DonorDNA | Sequence |
|---|---|
| SpE-R2toBam | CATCCTGCATGACCCCCCACCTCCCCAACCAGGAAGGCGGttttggatcctttttT GCCCCTGAATCCAGGAAAGCTGAGAATGCTTAGCTCCTT |

Genotyping PCR primers

| Primer name | Sequence | Amplicon |
|---|---|---|
| SpE-Fw | CATGTCTCTCTGTCTTGGGTGA | WT: 655 bp |
| SpE-Re | AGTTCATTCTGATGTCCCCAGT | Del: 213 bp |

***SpE-R2m mice give the same size (654-bp) of amplicon, but BamHI digestion of the amplicon gives 429-bp and 225-bp fragments.**

**Supplemental Table 1**. **Nucleotide sequences of guide RNA targets and PCR primers.** The nucleotide sequence of the targets for guide RNAs are shown in the upper table. Small characters represent the protospacer adjacent motif sequences. Nucleotide sequences of donor DNA used for R2 mutagenesis is shown in the middle table. Nucleotide sequences of PCR primers used for genotype determination are also shown in the lower table.
